# Migratory divides coincide with species barriers across replicated avian hybrid zones above the Tibetan Plateau

**DOI:** 10.1101/698597

**Authors:** Elizabeth S.C. Scordato, Chris C.R. Smith, Georgy A. Semenov, Yu Liu, Matthew R. Wilkins, Wei Liang, Alexander Rubtsov, Gomboobaatar Sundev, Kazuo Koyama, Sheela P. Turbek, Michael B. Wunder, Craig A. Stricker, Rebecca J. Safran

## Abstract

Migratory divides are proposed to be catalysts for speciation across a diversity of taxa. However, the relative contribution of migratory behavior to reproductive isolation is difficult to test. Comparing reproductive isolation in hybrid zones with and without migratory divides offers a rare opportunity to directly examine the contribution of divergent migratory behavior to reproductive barriers. We show that across replicate sampling transects of two pairs of barn swallow (*Hirundo rustica*) subspecies, strong reproductive isolation coincided with an apparent migratory divide spanning 20 degrees of latitude. A third subspecies pair exhibited no evidence for a migratory divide and hybridized extensively. Within migratory divides, migratory phenotype was associated with assortative mating, implicating a central contribution of divergent migratory behavior to reproductive barriers. The remarkable geographic coincidence between migratory divides and genetic breaks supports a longstanding hypothesis that the Tibetan Plateau is a substantial barrier contributing to the diversity of Siberian avifauna.

## Introduction

Migratory divides- regions where sympatric breeding populations overwinter in different geographic locations- have been proposed to facilitate completion of the speciation process by generating reproductive barriers that maintain species boundaries. Migratory divides can lead to prezygotic reproductive barriers via assortative mating if individuals with different wintering grounds arrive to breed at different times (Bearhop *et al*. 2005; Rolshausen *et al*. 2009; Taylor & Friesen 2017). They can also accelerate the evolution of postmating barriers if hybrids incur survival costs associated with the use of maladaptive routes between breeding and nonbreeding locations (Helbig 1991, 1996; Berthold *et al*. 1992; Delmore & Irwin 2014; Lundberg *et al*. 2017). However, establishing a clear link between divergent migratory behavior and reproductive isolation has been challenging. Migratory divides often occur at hybrid zones or regions of secondary contact, where evolutionary history, divergence in traits unrelated to migratory behavior, and ecological differences can also contribute to reproductive barriers (Ruegg 2008; Ruegg *et al*. 2012; Delmore *et al*. 2016; Toews *et al*. 2017). Isolating the effects of migratory behavior on reproductive barriers is particularly challenging when a single region of contact is examined between taxa with broad geographic distributions, because it is not possible to assess the generality of divergent migratory behavior in restricting gene flow across the species range. We therefore lack a comprehensive understanding of the relative importance of divergent migratory behavior to the formation and maintenance of species boundaries (Turbek *et al*. 2018).

Here we evaluate the hypothesis that migratory divides play a central role in the maintenance of reproductive isolation in secondary contact. We specifically examine three predictions of this hypothesis. First, hybridization should be more limited in contact zones with migratory divides compared to contact zones without migratory divides, when controlling for evolutionary history and divergence in non-migratory traits. Second, if migratory divides *per se* limit hybridization, migratory phenotype should explain a larger proportion of genetic variance among individuals than other divergent traits within migratory divides. Third, if migratory divides act as important premating reproductive barriers, then assortative mating by migratory phenotype should be stronger than assortative mating based on other divergent traits. Previous studies have found mixed evidence for assortative mating and genetic differentiation at migratory divides (Turbek *et al*. 2018), but have not assessed the relative contributions of different traits to reproductive barriers or compared reproductive isolation in hybrid zones with and without migratory divides. We evaluate these predictions in three subspecies of barn swallow (*Hirundo rustica*) that hybridize in Asia.

Barn swallows comprise six subspecies, of which three (*H. r. rustica, H. r. tytleri, and H. r. gutturalis*) are long-distance migrants that diverged in allopatry (Zink *et al*. 2006; Dor *et al*. 2010) but now share breeding range boundaries in Siberia and central Asia (Scordato & Safran 2014). There is a narrow hybrid zone in central Siberia between *rustica* and *tytleri,* but extensive hybridization in eastern Siberia between *tytleri* and *gutturalis* (Scordato *et al*. 2017, Figure 1). Differentiation in mtDNA is shallow and indicates that *gutturalis* and *tytleri* are more closely related to one another than either is to *rustica* (Zink *et al*. 2006; Dor *et al*. 2010), but genome- wide pairwise FST is similarly small (∼0.02) among allopatric populations of all three subspecies (Scordato *et al*. 2017). There is thus dramatic variation in the strength of reproductive isolation between subspecies, despite similarly shallow genetic differentiation.

**Figure 1:**
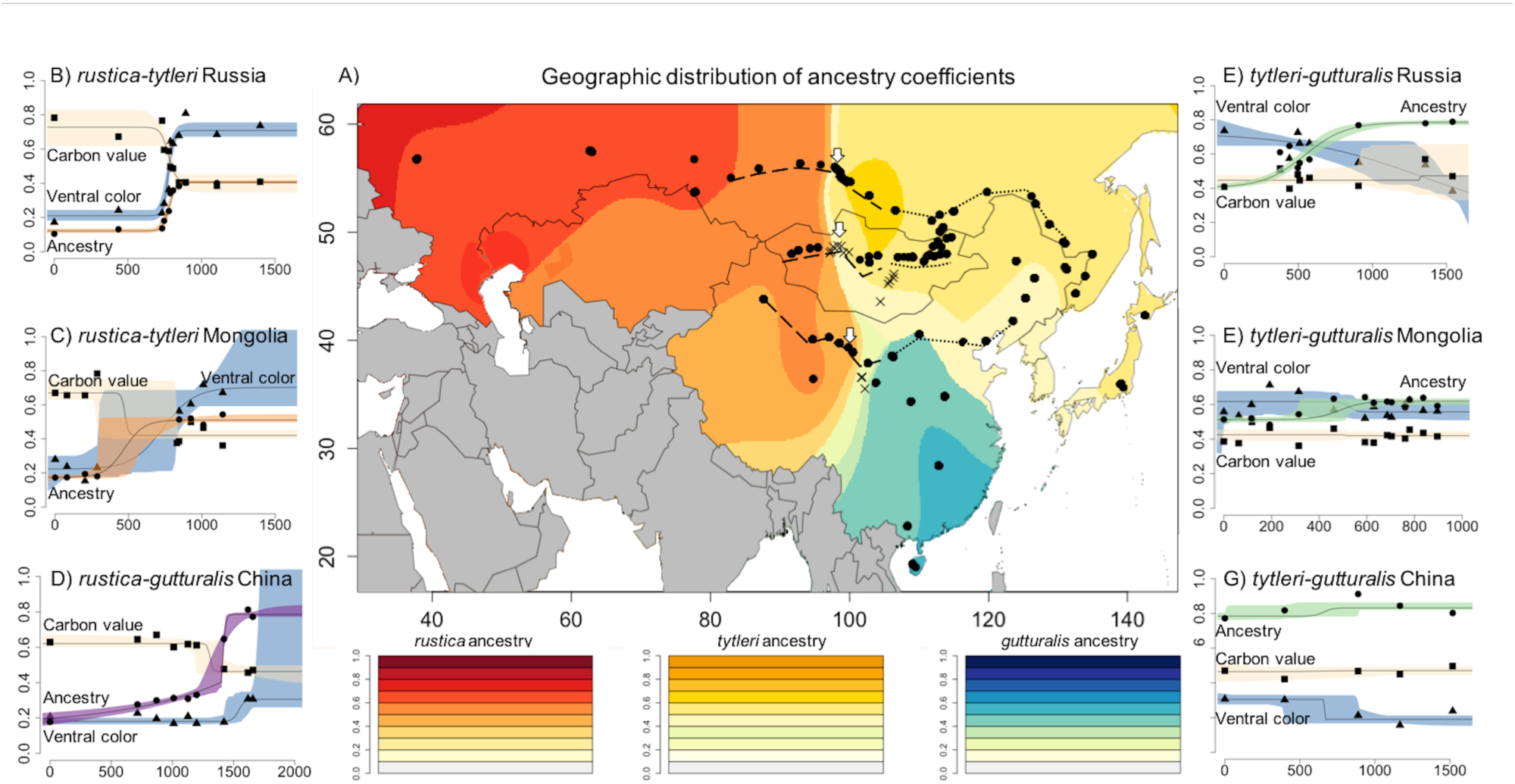
Geographic variation in ancestry coefficients and phenotypes across Asia. A) ancestry coefficients derived from spatially explicit modeling of >12,000 SNPs. Darker colors reflect more parental-like ancestry (red=*rustica,* gold=*tytleri,* blue*=gutturalis;* see legend). Paler colors reflect regions with more admixed individuals. Points indicate sampling locations. X’s are surveyed regions with no breeding barn swallows. Dashed lines show *rustica* transects used in geographic cline analysis (left panels) and dotted lines show *tytleri-gutturalis* transects (right panels). All cline plots show standardized trait values (y-axis) plotted against distance from the westernmost point of the transect (x-axis). **Left panels:** clines for genetic ancestry (orange: *rustica-tytleri*; purple: *rustica-gutturalis*), carbon value (tan clines), and ventral coloration (blue clines) across the three western sampling transects (B: Russia; C: Mongolia; D: China). Clines for carbon value and ancestry are steep and coincident across all three contact zones, and cline centers all occur at 98-100 degrees longitude (centers marked on map with white arrows). **Right panels:** geographic clines for ancestry (green clines: *tytleri-gutturalis*), carbon value (tan clines), and ventral coloration (blue clines) across the three eastern sampling transects (E: Russia; F: Mongolia; G: China). Ancestry clines are shallow and broad, and there is no variation in isotope values and little variation in ventral color across the transects.

We evaluated the extent to which a migratory divide explains this variation in strength of reproductive isolation. The geographic location of the narrow hybrid zone in Siberia coincides with reported migratory divides in several other pairs of avian taxa (Irwin & Irwin 2005). The convergence of migratory divides in this region may be caused by the Tibetan Plateau: small-bodied passerines tend to migrate to the west or east around this geographic barrier (Irwin & Irwin 2005). Divergent migratory behavior has therefore been proposed to be broadly important to the evolution and maintenance of species boundaries in Siberian avifauna (Irwin & Irwin 2005). However, barn swallow subspecies also differ in ventral plumage coloration, tail streamer length, and body size (Turner 2010; Scordato & Safran 2014), and these traits could be more important reproductive barriers than migratory behavior. We quantified the relative contribution of migratory behavior to reproductive barriers via comprehensive measurement of phenotype, detailed genomic analyses, and measures of assortative mating. We applied these measures to replicated transects to assess the generality of our results across a large proportion of the species range.

## Materials and Methods

### Sampling

We sampled 1288 birds across the range boundaries of the three Eurasian barn swallow subspecies (Figures 1, 2). In addition to previously sampled hybrid zones between *rustica-tytleri* and *tytleri-gutturalis* in Russia (Scordato *et al*. 2017), we discovered a hybrid zone between *rustica* and *gutturalis* in western China, as well as additional regions of contact between *tytleri*- *gutturalis* and *rustica-tytleri* in Mongolia and China (Figures 1, 2).

**Figure 2:**
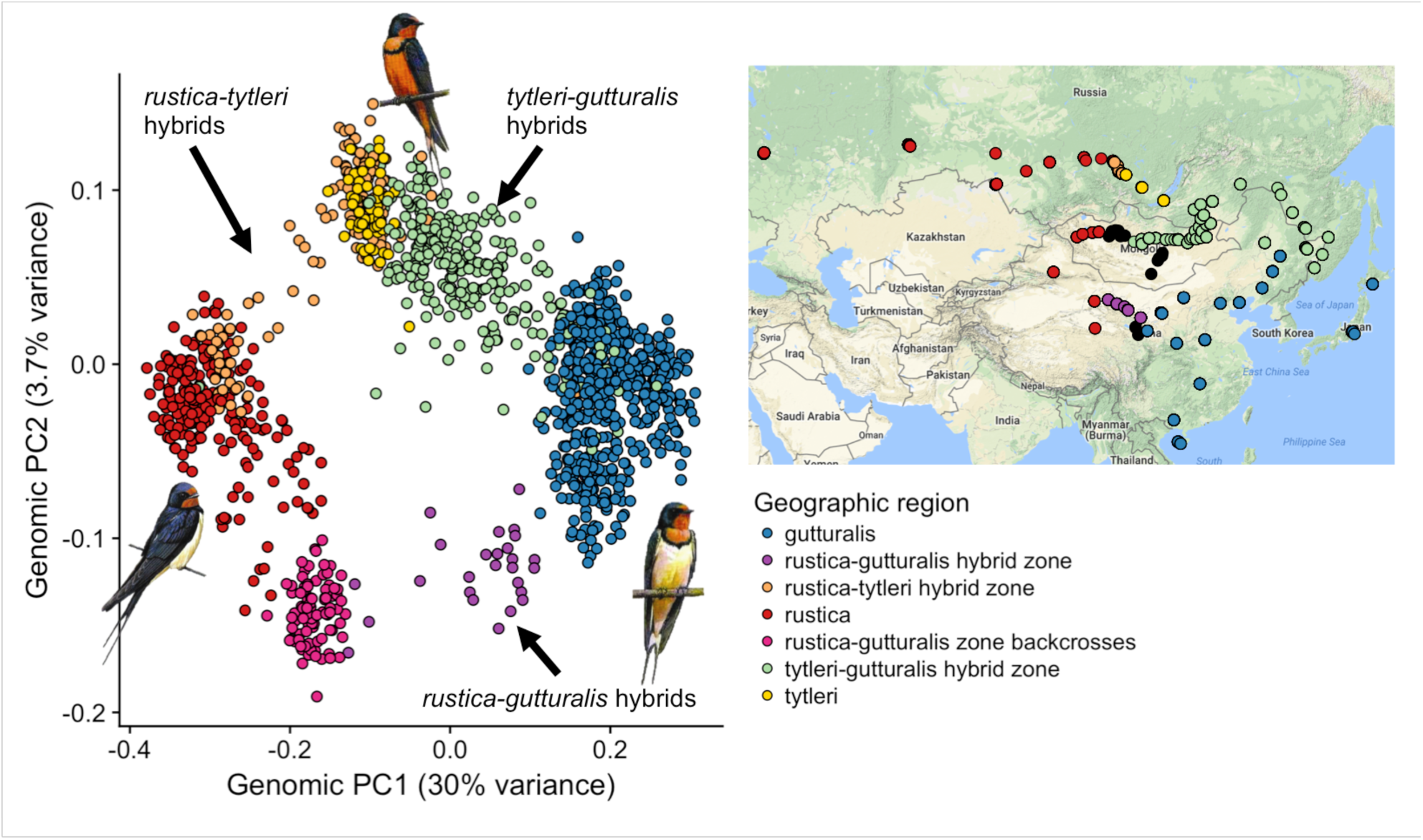
The first two principal components from a PCA of the genetic covariance matrix of all individual birds. Point colors correspond to geographic sampling regions, as indicated in the legend and on the inset map. The PCA generally recapitulates geography and recovers three parental clusters (*rustica* in red, *tytleri* in gold, and *gutturalis* in blue). Points connecting these clusters correspond to admixed individuals (labeled arrows). The pink cluster at PC1= −0.2 are birds captured in the *rustica-gutturalis* hybrid zone (purple points on map) that appear to be late generation backcrosses to *rustica* and form their own discrete genetic cluster. Drawings show typical phenotype for each of the three parental subspecies. Inset: map of sampling locations with points colored corresponding to geography in the main figure legend. Black points are sampled areas where no birds were found to be breeding. Drawings courtesy of Hilary Burns.

Sampling was conducted during barn swallow breeding seasons (April-July 2013 in Russia, April-July 2014 in China, Mongolia, and Japan, and May-June 2015 in western China). Birds were caught in mist nets and individually banded with numbered aluminum leg bands. An ∼80ul blood sample was collected via brachial venipuncture and stored in Queen’s lysis buffer. We collected 5-10 feathers from the throat, breast, belly, and vent of each bird for quantification of color, and collected the inner two tail rectrices for analysis of stable isotopes. Length of the right wing chord, tail streamers, and each primary feather were measured to the nearest 0.1mm, and weight was measured to 0.5g. Each morphometric measurement was taken 3 times per bird, and averaged measurements were used in subsequent analyses. The length of the primary feathers was used to calculate wing pointedness and convexity (Lockwood *et al*. 1998). Wing length has been used as a proxy for migratory distance (Safran *et al*. 2016), but is also correlated with body size, whereas wing shape (pointedness and convexity) has been explicitly linked to migratory distance and is independent of body size (Lockwood *et al*. 1998).

### Social pair identification

Barn swallows are socially monogamous, with both males and females building the nest and provisioning offspring (Turner 2010). To assess assortative mating, we assigned birds to a social pair if the male and female were unambiguously caught at the same nest. It was not possible to assign birds to pairs in large colonies because they were not caught at individual nests. Our measures of assortative mating are therefore derived from birds nesting singly or in small groups.

### Quantification of color, identification of variants

We analyzed plumage color using a spectrophotometer. DNA was extracted and sequenced on four replicate Illumina HiSeq lanes. Reads were aligned to a draft barn swallow reference genome (Safran *et al*. 2016) and variants called using *bcftools* and *samtools* (Li & Durbin 2009; Li *et al*. 2009). We identified 12,383 single nucleotide polymorphisms (SNPs) with 5% minor allele frequency cutoff and median read depth of seven reads per locus. Methods for color quantification, sequencing, and variant calling are described elsewhere (Safran & McGraw 2004; Hubbard *et al*. 2015; Safran *et al*. 2016; Scordato *et al*. 2017; Smith *et al*. 2018) and are detailed in the Supplemental Material.

## Analysis

### Evidence for a migratory divide

We assessed evidence for migratory divides by analyzing stable carbon (δ^13^C) values in tail feathers collected from birds on the breeding grounds (see Supplemental Material). Barn swallows molt their tail feathers in the winter (Turner 2010). Because feather keratin is metabolically inert after formation, feathers sampled during the summer reflect isotopic environments occupied during winter, when feathers were grown. Stable isotope values do not provide direct information about geographic locations of feather growth. However, environmental δ^13^C values vary systematically and widely with water use efficiency of plants; this differentiation is preserved through the food web to animals, such that large differences in feather δ^13^C between individuals suggest those individuals grew their feathers in different environments (Kelly 2000). We evaluated differences in the distribution of δ^13^C values between each of the three subspecies and among hybrids in regions of secondary contact. We found support for migratory divides between *rustica-tytleri* and *rustica-gutturalis* (see Results, Figures 1, 2). We use δ^13^C values (hereafter “carbon isotope values”) as proxies for an individual’s migratory phenotype in subsequent analysis.

### Prediction one: population structure and extent of hybridization

#### Population structure

We used three complementary methods to analyze population structure: principal components analysis (PCA), which does not require an *a priori* number of populations; TESS (Caye *et al*. 2016), a spatially explicit clustering method that assigns individuals to *K* clusters but weights individual admixture proportions by geographic proximity; and fastSTRUCTURE (Raj *et al*. 2014), which uses a variational Bayesian algorithm to assign individuals to *K* clusters without weighting by geographic proximity. We ran the PCA on the genome-wide covariance matrix of 12,383 SNPs across 1288 individuals using the R function *prcomp*. We ran TESS on the same set of SNPs for values of K from 2-5, with 3 repetitions per K, 1000 iterations, and the regularization parameter (alpha)= 0.001. This regularization value does not weight geographic location particularly strongly in the analysis (Caye *et al*. 2016). We ran the fastSTRUCTURE model with the “simple” prior for values of K from 1-15 and a cross-validation of 5 repetitions per K. In fastSTRUCTURE, the best value of K is the minimum number of model components (K) that explain 99.99% of the admixture in the sample.(Raj *et al*. 2014) We found K=3 to be the best value. We assigned individual birds to hybrid classes (F1, later generation hybrid, or backcross) by calculating hybrid indices and average heterozygosity across subsets of differentiated loci using the R package *introgress* (Supplemental Material).

#### Geographic cline analysis

To determine whether geographic variation in the frequency of hybridization coincides with differences in migratory behavior or other divergent phenotypic traits, we fit sigmoidal geographic clines (Szymura & Barton 1986) to three east-west transects spanning contact zones between *rustica*-*tytleri* (two transects) and *rustica-gutturalis* (one transect, Figure 1). Transects spanned 85-115 degrees longitude. We explicitly compared the extent of hybridization in regions with and without putative migratory divides by fitting clines to three parallel transects at the same latitudes but farther-eastern longitudes (106-140 degrees) through regions of admixture between *tytleri* and *gutturalis* (Figure 1). Clines were fit to genomic ancestry, measured as PC1 from the PCA of the genome-wide covariance matrix. PC1 explained 30% of the genetic variance and clearly separated the three subspecies as well as hybrids (Figure 2). To assess whether variation in phenotype was geographically concordant with admixture, we also fit clines to breast chroma, throat chroma, carbon isotope value, tail streamer length, wing convexity, wing pointedness, and wing length. Cline analysis was implemented in the R package HZAR (Derryberry *et al*. 2014, Supplemental Material). We applied neutral diffusion equations (Barton & Gale 1993) to determine whether cline widths were narrower than expected under a scenario of no selection or reproductive isolation, assuming a one-year generation time and dispersal distances of 42km (conservative) or 100km (less conservative, Paradis *et al*. 1998; Supplemental Material). Cline widths narrower than the neutral expectation may be maintained by selection and contribute to reproductive barriers (Ruegg 2008; Brelsford & Irwin 2009). Concordant clines between ancestry and phenotypic traits may indicate that those traits are associated with reproductive barriers (Gay *et al*. 2008; Gompert & Buerkle 2016).

### Prediction two: variance partitioning

To test the prediction that traits associated with reproductive barriers explain comparatively large proportions of genetic variance, we partitioned genetic variance among groups of traits using variance partitioning and redundancy analysis in the *ecodist* and *vegan* packages in R (Goslee & Urban 2007; Oksanen *et al*. 2013). This approach determines the amount of variance in a set of response variables that is due to a set of explanatory variables, while conditioning on other sets of variables. It is ideal for large datasets with intercorrelated explanatory variables (Wang 2013; Safran *et al*. 2016). We quantified the amount of variance in genomic PC1 and PC2 (Figure 2) that could be explained by the individual and combined contributions of migratory phenotype (carbon isotope value) and ventral coloration. The broad geographic scale of sampling required controlling for possible isolation-by-distance (Shafer & Wolf 2013; Wang 2013). We therefore analyzed each transect separately and conditioned models on sampling location (latitude and longitude).

### Prediction three: assortative mating

Premating reproductive isolation is maintained by assortative mating between individuals with similar genotypes (“like mating with like). However, premating isolation is typically measured by assessing assortative mating by phenotype, under the assumption that phenotype is a reasonable proxy for genotype. Interpreting assortative mating is complicated when there is continuous variation in phenotypes and genotypes between interbreeding groups. We therefore measured assortative mating in two ways. First, we used phenotype networks to identify correlations between an individual’s genotype and its mate’s phenotype. This method leverages continuous variation in genotypes and phenotypes to quantify broad patterns of assortative mating across sampling transects. Second, we calculated standardized indices of reproductive isolation within populations to determine the strength of assortative mating based on different traits (genotype, migratory phenotype, and ventral color). These two methods provide complementary views of assortative mating at different geographic scales. We were able to assign birds to social pairs along three sampling transects: the *rustica-tytleri* transect in Russia, the *rustica-gutturalis* transect in China, and the *tytleri-gutturalis* transect in China (Figure 1). Sufficient social pairing data were not available for the other three transects.

### Assortative mating: phenotype networks

To accommodate continuous variation between parentals and hybrids we used a Partial Correlation and Information Theory (PCIT) approach (Badyaev & Young 2004; Wilkins *et al*. 2015) to identify correlations between male and female phenotypes and genotypes. This method was originally developed for analysis of gene co-expression networks (Reverter & Chan 2008) but is applicable to other networks with complex correlation structures (Shizuka & Farine 2016). We began with a matrix of Spearman rank correlations between pairs of males and females. These matrices included genotype (genomic PC1), ventral color, carbon isotope value, and sampling latitude and longitude for each member of a social pair. To identify and remove spurious correlations, we used the *pcit* package in R (Watson-Haigh *et al*. 2009), which uses the Spearman rank correlation matrix to generate a network of partial correlation coefficients. The PCIT algorithm sets a ‘local threshold’ for inclusion of an edge (i.e. the correlation connecting two traits) based on the average ratio of the partial to direct correlation for every trio of traits (“nodes” on the network). The algorithm begins with a network in which every pair of nodes is connected by an edge whose value is the absolute value of the correlation coefficient between the two traits. An edge between two particular nodes is discarded if the direct correlation coefficient is less than the product of the local threshold and the correlations between each node in the focal pair and the third trait in the trio.

We visualized assortative mating for each transect as a bipartite network of correlations with two categories of nodes (male and female). Each node represents a different trait, and lines (edges) connect nodes if traits are correlated within mated pairs (e.g. if darker males mate with darker females; Figure 5, gray lines). Analyzing assortative mating along the transects ensured that each network encompassed individuals with parental and admixed genotypes. Including genotype as a node in the network allowed us to determine which aspects of phenotype might be used as reliable proxies of genotype in the context of maintaining subspecies boundaries. These relationships are shown as black lines in Figure 5 connecting an individual’s genotype to the phenotype of its social partner. We generated networks using the R package ‘qgraph’(Epskamp *et al*. 2012). To facilitate interpretation, we only show correlations between male and female pairs on the networks (as opposed to within-individual trait correlations), but within-individual correlations were included in the PCIT analysis.

### Assortative mating: strength of premating isolation

To examine fine-scale assortative mating within populations, we analyzed the strength of premating reproductive isolation (RI) following Sobel and Chen (2014). Here, isolation is calculated based on the proportion of heterospecific pairings divided by the sum of conspecific and heterospecific pairings. This method is advantageous because RI is scaled between −1 and 1, with 1 equal to complete assortative mating, 0 equal to random mating, and −1 equal to complete disassortative mating. The isolation index is directly related to gene flow: RI = 0.5 means there are 50% fewer heterospecific pairs in the population than expected by chance, whereas RI = −0.5 means there are 50% more heterospecific pairs than expected by chance.

This RI index requires assigning individuals to categories to determine frequencies of con- vs. heterospecific pairings. We assigned each individual as a “parental” or a “hybrid” based on its genotype, its migratory phenotype, and its color. Assignments were made using 1000 repetitions of a linear discriminant analysis (see Supplemental Material). We then calculated the strength of RI based on each trait in each population across the three transects.

Because genotype frequencies (i.e. the proportions of parentals vs. hybrids) varied between populations, we followed equation 4S4 in Sobel and Chen (2014) and weighted observed con- and heterospecific pairings by the number of such pairings expected under a scenario of random mating, given the distribution of genotypes in the population:

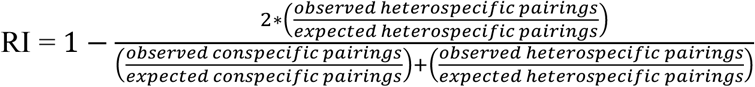

To calculate expected pairings, we used the total pool of individuals (not just those for which we had pairing data) and randomly generated social pairs without replacement. We counted the proportions of con and heterospecific pairs from these random draws. We considered pairings between two hybrid individuals to be “conspecific” and pairings between a parental and a hybrid to be “heterospecific;” this will generally underestimate the strength of reproductive isolation. The expected proportions of each type of pairing under a random mating scenario were averaged over 1000 random draws for each population.

## Results

### Evidence for a migratory divide

The distribution of δ^13^C in feathers for *tytleri* overlapped almost completely with *gutturalis*, whereas the distribution for *rustica* minimally overlapped the distributions for the other two subspecies (Figure 4, Figure S1). More importantly, the δ^13^C values for *rustica* are consistent with comparatively arid environments where food webs are based on C4 plants, whereas the values for *gutturalis* and *tytleri* are consistent with more mesic environments where food webs are based on C3 plants (Kelly 2000). Furthermore, observed δ^13^C values for *rustica* are consistent with values expected for southern and eastern Africa and the Arabian peninsula, a region dominated by C4 plants (Still *et al*. 2003), and an area of extensive sighting records (Sullivan *et al*. 2009; Turner 2010) for this subspecies during winter. By contrast, both δ^13^C distributions and sighting records suggest *tytleri* and *gutturalis* overwinter in south and southeast Asia, a wetter region with comparatively more C3 plants (Still *et al*. 2003). Hybrid zones between *rustica* and *tytleri/gutturalis* exhibit intermediate means and large variances in δ^13^C values (Figure S1), suggesting sympatry between individuals overwintering in different locations. We interpret these results as evidence for different wintering grounds and consequent migratory divides between *rustica*-*tytleri* and *rustica-gutturalis* (Figure 1).

### Prediction 1: Limited hybridization is associated with divergent migratory phenotypes

We predicted that if migratory divides act as barriers to reproduction, then hybridization should be limited in contact zones with migratory divides compared to contact zones without migratory divides. Furthermore, clines for carbon isotope values, our proxy for differences in overwintering grounds, should be steep and concordant with genetic ancestry clines.

#### Population structure and gene flow

We identified three genetic clusters corresponding to the three subspecies, with dramatic variation in the extent of hybridization between subspecies pairs (Figure 1,2). We found narrow hybrid zones between *rustica-tytleri* and *rustica-gutturalis,* whereas *tytleri* and *gutturalis* were admixed over a large region of east Asia (Figures 1, 2).

We found F1, later generation hybrid, and backcrossed individuals between all three subspecies pairs, indicating ongoing gene flow (Figure S2). However, there were few recent hybrids between *rustica-tytleri* (1% F1, 13% later generation) and *rustica*-*gutturalis* (2% F1, 18% later generation, Figure S2), consistent with strong isolation between these two subspecies pairs. By contrast, there were many multi-generation hybrids between *tytleri* and *gutturalis* (8% F1 and 53% later generation; Figure S2), consistent with weak reproductive isolation across a broad geographic region that contains few parental individuals and many hybrids. These analyses reveal less hybridization overall between the subspecies pairs with migratory divides (*rustica- tytleri, rustica-gutturalis*) compared to the pair without a migratory divide (*tytleri-gutturalis*).

#### Geographic clines- rustica pairs

Clines for genetic ancestry (genetic PC1) were very narrow between *rustica-tytleri* in Russia and *rustica-gutturalis* in China, suggesting these hybrid zones are maintained by selection or are of unrealistically recent origin (<1 year; Figure 1B, D, Table 1). A mountain range separated *rustica* and *tytleri* in western Mongolia, and we found no evidence for extant interbreeding across this barrier (Figure 1C, Table 1). Remarkably, the centers of the ancestry clines in all three *rustica* transects occurred at similar longitudes (between 98 and 101 degrees), despite spanning over 20 degrees of latitude and comprising different pairs of subspecies (Figure 1A, white arrows). Carbon isotope clines were narrow and concordant with ancestry in all three *rustica* transects (Figure 1, Table 1). The locations of narrow hybrid zones thus coincide with migratory divides.

**Table 1:**
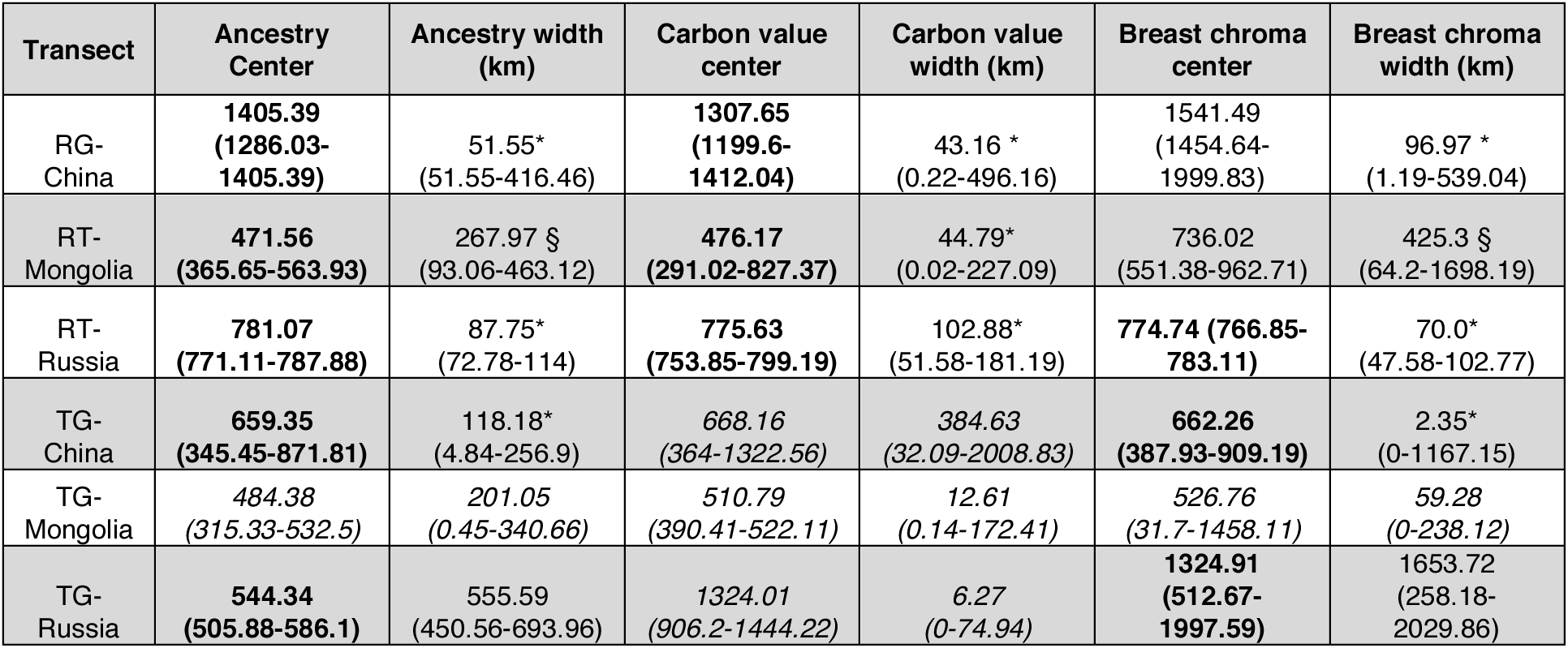
Best-fit geographic cline models for each trait and each transect. Boldfaced clines are those that have centers coincident with the ancestry cline. Starred widths are narrower than expected under a neutral diffusion model assuming a dispersal distance of 42 km and a hybrid zone age older than 20 years. The § symbol shows clines that are wider than expected with a dispersal distance of 42km, but narrower than expected if dispersal is 100km and clines are older than 20 years. Italicized clines show no statistically significant variation in trait values across the transect and are consequently poorly described by cline models. Carbon clines coincide with ancestry in the three *rustica* transects (top three rows) but not in the *tytleri-gutturalis* transects (bottom three rows). Cline center units are kilometers from the westernmost transect point.

| Transect | Ancestry Center | Ancestry width (km) | Carbon value center | Carbon value width (km) | Breast chroma center | Breast chroma width (km) |
| --- | --- | --- | --- | --- | --- | --- |
| RG-China | **1405.39**<br>(**1286.03-1405.39**) | 51.55*<br>(51.55-416.46) | **1307.65**<br>(**1199.6-1412.04**) | 43.16 *<br>(0.22-496.16) | 1541.49<br>(1454.64-1999.83) | 96.97 *<br>(1.19-539.04) |
| RT-Mongolia | **471.56**<br>(**365.65-563.93**) | 267.97 §<br>(93.06-463.12) | **476.17**<br>(**291.02-827.37**) | 44.79*<br>(0.02-227.09) | 736.02<br>(551.38-962.71) | 425.3 §<br>(64.2-1698.19) |
| RT-Russia | **781.07**<br>(**771.11-787.88**) | 87.75*<br>(72.78-114) | **775.63**<br>(**753.85-799.19**) | 102.88*<br>(51.58-181.19) | **774.74 (766.85-783.11)** | 70.0*<br>(47.58-102.77) |
| TG-China | **659.35**<br>(**345.45-871.81**) | 118.18*<br>(4.84-256.9) | *668.16*<br>(*364-1322.56*) | *384.63*<br>(*32.09-2008.83*) | **662.26**<br>(**387.93-909.19**) | 2.35*<br>(0-1167.15) |
| TG-Mongolia | *484.38*<br>(*315.33-532.5*) | *201.05*<br>(*0.45-340.66*) | *510.79*<br>(*390.41-522.11*) | *12.61*<br>(*0.14-172.41*) | *526.76*<br>(*31.7-1458.11*) | *59.28*<br>(*0-238.12*) |
| TG-Russia | **544.34**<br>(**505.88-586.1**) | 555.59<br>(450.56-693.96) | *1324.01*<br>(*906.2-1444.22*) | 6.27<br>(0-74.94) | **1324.91**<br>(**512.67-1997.59**) | 1653.72<br>(258.18-2029.86) |

Ventral coloration also varied among *rustica* pairs. A narrow ventral color cline in Russia coincided with the ancestry and carbon isotope clines (Figure 1, Table 1). Ventral coloration differed on either side of the mountains in Mongolia (Table 1). In the *rustica*-*gutturalis* transect in China, the cline for color was narrow but the center was displaced to the east of the other clines (Figure 1, S1, Table 1). There may thus be some differential introgression of plumage color between *rustica* and *gutturalis,* although differences in color were small because both subspecies have mostly white ventral plumage (Figure S1).

Clines for wing pointedness were narrow and coincident with the ancestry and carbon isotope clines in the two *rustica-tytleri* transects, but did not vary across the *rustica-gutturalis* transect (Table S1). Tail streamer length, throat color, wing convexity, and wing length either did not vary clinally or exhibited very wide clines (Table S1). Thus, carbon isotope value (reflecting different wintering grounds) was the only trait consistently associated with genetic ancestry and limited hybridization across the *rustica* pairs. This result supports our prediction that narrow hybrid zones are associated with migratory divides. The convergent geographic locations of ancestry and migratory clines strongly suggests that differences in wintering grounds are driven by divergent migratory routes around the Tibetan Plateau (Figure 1).

#### Geographic clines- tytleri/gutturalis pair

There was extensive admixture and no clear association between genetic ancestry and phenotype across the three *tytleri –gutturalis* transects. Ancestry clines were wide, with only the cline in China narrower than the neutral expectation (Table 1, Figure 1 E-F). Furthermore, there was no clinal variation in ancestry across Mongolia, indicative of homogenous admixture (Figure 1F, S1). There was also no clinal variation in carbon isotope values across any of the three *tytleri-gutturalis* transects (Figure 1, Table 1). The only transect with a ventral color cline narrower than the neutral expectation was in China, where the cline was concordant with ancestry (Table 1). As with the *rustica* pairs, morphometric traits did not vary clinally across the transects (Table S1). These analyses reveal large geographic regions of nearly homogenous admixture and little phenotypic differentiation between *tytleri* and *gutturalis*, in contrast to the narrow hybrid zones that coincided with migratory phenotype and, to some extent, color, in the *rustica* migratory divides.

### Prediction 2: Migratory phenotype is associated with genetic differentiation

We predicted that if migratory divides are important reproductive barriers, differences in migratory behavior would explain large proportions of among-individual genome-wide variance relative to other divergent traits within migratory divides. The combined effects of color, migratory phenotype, and geographic location explained 34% of among-individual genetic variance (PC1 and PC2) in the Russian *rustica-tytleri* transect (Figure 3A). The combination of geography and migratory phenotype explained 30% of genetic variance between *rustica-tytleri* in Mongolia (Figure 3C) and 23% between *rustica*-*gutturalis* in China (Figure 3F). Migratory phenotype explained statistically significant proportions of genetic variance when controlling for the effects of color and geography in the *rustica-tytleri* transect in Russia and the *rustica- gutturalis* transect in China (Table S2). Color explained significant proportions of genetic variance in the two *rustica-tytleri* transects when controlling for geography and migratory phenotype, but not in the *rustica-gutturalis* transect in China (Table S2). Overall, the combination of migratory phenotype and geography explained larger proportions of variance than did geography and color in two of the three *rustica* transects. The combination of all three factors explained the largest proportion of variance in the *rustica-tytleri* transect in Siberia (Figure 3).

**Figure 3:**
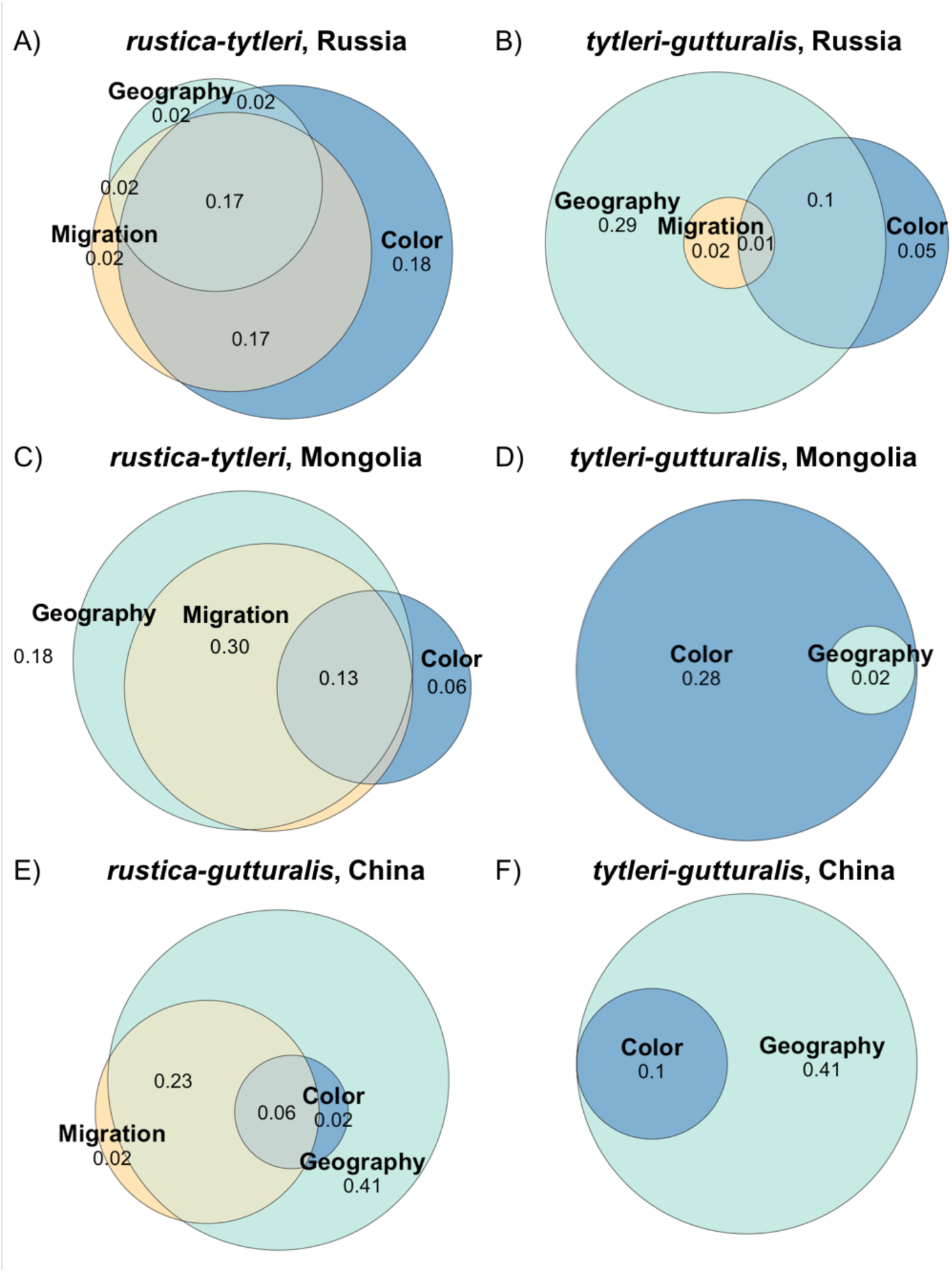
Genomic variance (PC1 and PC2) partitioned among traits related to migratory phenotype (carbon value), sexual signaling (ventral color), and geographic location of sampling (latitude and longitude). Variance is shown as adjusted R^2^ values. Variance is partitioned among the subsets of individuals occurring along each of the six transects through regions of hybridization shown in Figure 1. Each row shows a transect with a migratory divide on the left and the parallel transect (same latitude, different longitude) without a migratory divide on the right. Overlapping regions between circles show the amount of genetic variance explained by the combined effects of those variables; for example, the combination of migratory phenotype, color, and geographic location explains 17% of the genetic variance in the *rustica-tytleri* transect in Russia (A) and the combination of migratory phenotype and geographic location explains 30% of the genetic variance in the *rustica-tytleri* transect in Mongolia (C). Note that migratory phenotype explains no genetic variance in the *tytleri-gutturalis* transects in Mongolia and China.

In the three *tytleri-gutturalis* transects without migratory divides, migratory phenotype explained a maximum of 2% of among-individual genetic variance when combined with geography (Figure 3 B, D, F, Table S2), consistent with no clear migratory divide in these regions. Geography and color explained comparatively larger proportions of genetic variance (2- 28%, Figure 3, Table S2).

We visualized these individual-level associations by plotting frequency distributions of genotypes and phenotypes. The two subspecies pairs with migratory divides exhibited bimodal distributions of carbon isotope values coinciding with bimodal distributions of genotypes, with the rare hybrids expressing trait values that spanned the full parental range (Figure 4A, B). By contrast, carbon isotope distributions were unimodal between all *tytleri* and *gutturalis* populations, and hybrid genotypes were common (Figure 4C). Distributions of ventral color showed a different pattern: *rustica* and *gutturalis* have white ventral plumage, whereas *tytleri* is dark brown, resulting in bimodal color distributions between *rustica-tytleri* and *tytleri-gutturalis* (Fig 4E, F) and a unimodal distribution between *rustica-gutturalis* (Fig 4D). Color distributions did not match genotype distributions: there was limited hybridization between *rustica-gutturalis* despite similar ventral color, and extensive hybridization between *tytleri-gutturalis* despite different ventral color.

**Figure 4:**
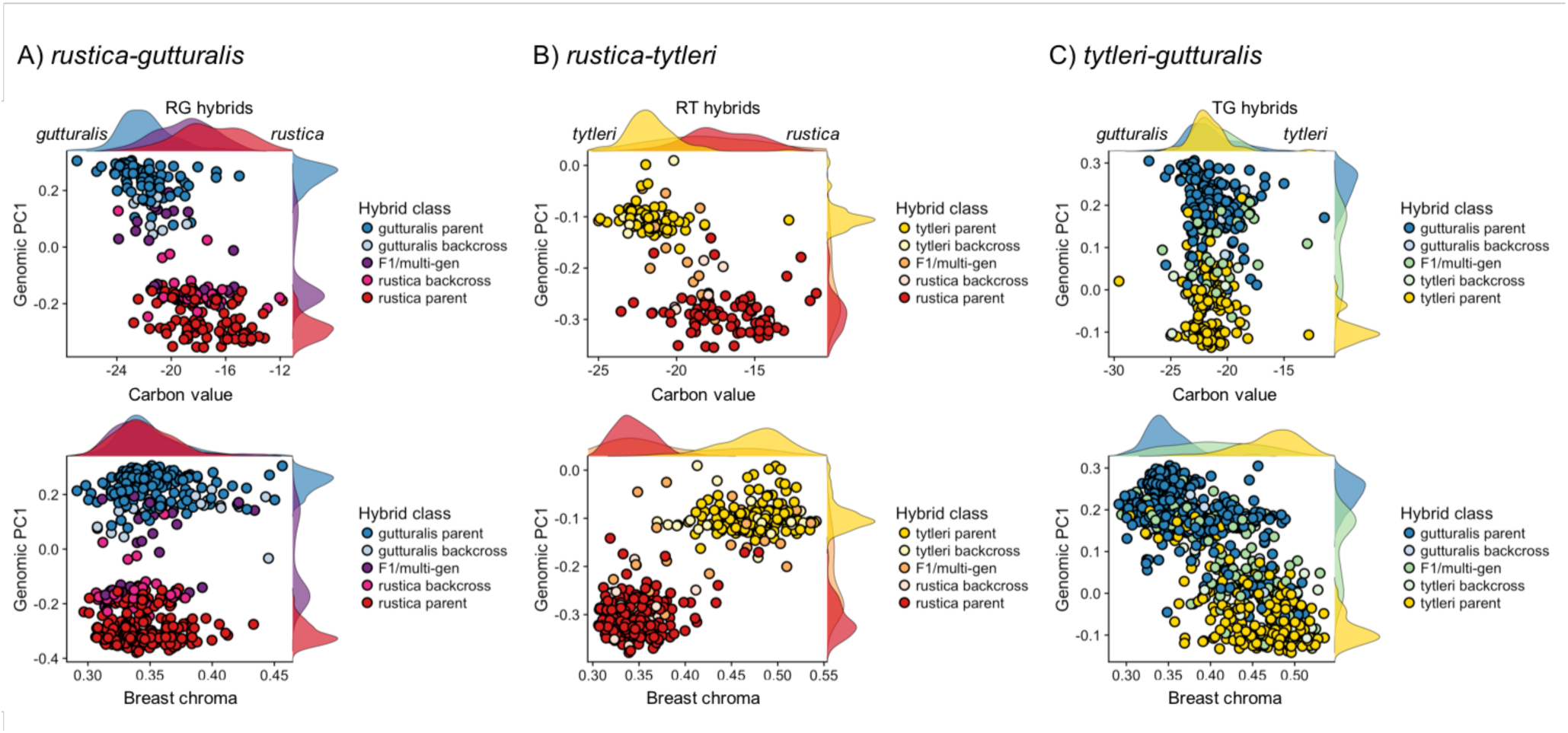
Distribution of genotypes and phenotypes between each of the three subspecies pairs. Each point is an individual bird, with points colored by hybrid class assignment (parental, backcross, or F1/multigenerational hybrid). Genomic PC1 score is on the y-axis of all plots. Y- axis density plots show the distribution of genomic ancestry for parentals and hybrids (backcrosses, F1, multigenerational hybrids combined) between each subspecies pair (red=*rustica*, blue=*gutturalis*, yellow=*tytleri*). Note clear bimodal distributions with few intermediates between *rustica-gutturalis* (purple) and *rustica-tytleri* (orange), but a broad range of genomically intermediate individuals between *tytleri* and *gutturalis* (green). There is also a separate peak in genomic PC scores in the rustica-gutturalis contact zone (A, purple peak on y- axis) comprised of late generation back-crossed individuals. **Top row**: Genomic PC1 score is plotted against carbon value(x-axis). There were generally bimodal distributions in carbon values between parental individuals in the *rustica-gutturalis* and *rustica-tytleri* pairs, corresponding to bimodal distributions of parental genotypes. In the *rustica-gutturalis* hybrid zone (A, purple), hybrid genotypes and carbon isotope ratios were more similar to parental *rustica*. In the *rustica-tytleri* hybrid zone (B, orange), hybrid genotypes and carbon isotope ratios were more similar to *tytleri*, but F1/multigenerational hybrids had isotope ratios spanning the full parental range. In the *tytleri-gutturalis* hybrid zones (C, green), there were no differences in isotope ratios between parentals and hybrids **Bottom row:** Genomic PC score is plotted against breast chroma (x-axis). A)There are few hybrids between *rustica* and *gutturalis* despite similar parental ventral color. B) In the *rustica-tytleri* and C) *tytleri-gutturalis* zones, we find bimodal distributions in ventral color. There are few hybrids between *rustica-tytleri* (B, orange) but many hybrids between *tytleri-gutturalis* (C, green), despite differences in ventral color.

### Prediction 3: Assortative mating is based on migratory phenotype

We found that divergent migratory phenotypes, and, to a lesser extent, divergent color were associated with limited hybridization and comparatively large genome-wide variance. Lastly, we predicted that if migratory behavior *per se* acts as a barrier to reproduction, we would observe assortative mating by migratory phenotype in hybrid zones with migratory divides. We assessed the contribution of migratory phenotype to premating reproductive isolation using social pairing data across three transects: the *rustica-tytleri* hybrid zone in Russia, the *rustica-gutturalis* hybrid zone in China, and the *tytleri-gutturalis* transect in China (Figure 1). The first two transects have migratory divides, while the third does not.

#### Phenotype networks

Phenotype networks indicated assortative mating by genotype across all three transects (Figure 5: black lines connecting male and female genotypes; *rustica-gutturalis* rgenotype = 0.82; *rustica-tytleri* rgenotype = 0.48; *tytleri-gutturalis* rgenotype = 0.50). In the two transects with migratory divides, carbon isotope values were correlated within pairs (Figure 5A, B, gray lines). An individual’s genotype also correlated with its mate’s carbon isotope value (Figure 5A, B, black lines; *rustica-gutturalis:* rcarbon = 0.56 and 0.36; *rustica-tytleri:* rcarbon = 0.56 and 0.47), suggesting that overwintering grounds are an important basis for assortative mating. Carbon values were not associated with assortative mating in the transect without a migratory divide (*tytleri-gutturalis,* Figure 5C).

**Figure 5:**
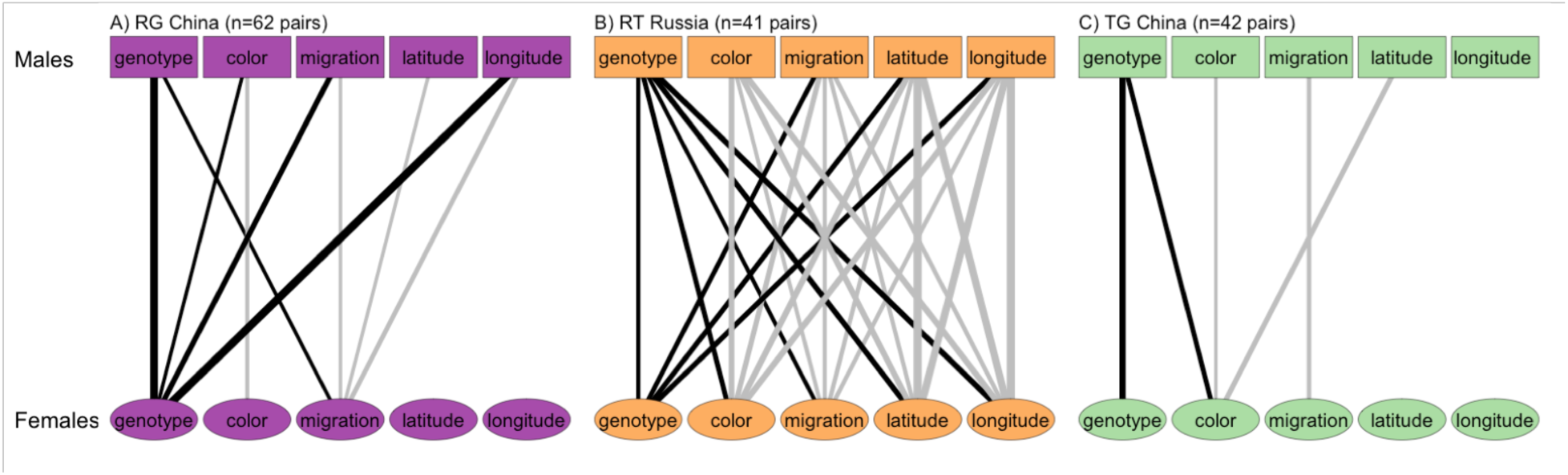
Bipartite phenotype networks showing traits associated with assortative mating across three transects. Black lines show characteristics that predict the ancestry of an individual’s social mate (the traits most relevant for reducing gene flow). Gray lines show traits correlated within pairs. Line width reflects the strength of the correlation. Squares (top row) are males. Circles (bottom row) are females. In the two migratory divides (A and B), an individual’s genotype is correlated with the migratory phenotype (carbon value) and ventral coloration of its social mate (black lines between traits and genotypes). There is also strong assortative mating by genotype. In the *tytleri-gutturalis* transect in China (C), there is assortative mating by genotype, and ventral coloration is associated with social mate’s genotype. In the migratory divides (A and B) there are also strong correlations between geographic location and genotype, indicating geographic variation in the distribution of available mates.

Ventral coloration was correlated with mate’s genotype in all three transects (*rustica- gutturalis*: rcolor = 0.35, *rustica-tytleri*: rcolor = 0.58, *tytleri-gutturalis*: rcolor = 0.38, Figure 4). The correlations for color were weaker than those for carbon isotopes in the *rustica-gutturalis* transect (Figure 5A) and similar in the *rustica-tytleri* transect (Figure 5B). We interpret this as evidence for migratory behavior and, to a lesser extent, coloration, in mediating broad patterns of assortative mating across hybrid zones with migratory divides However, genotype and phenotype also correlated with geographic location in all three transects (Figure 5). These correlations reflect geographic variation in the frequencies of different genotypes and phenotypes (Figure 1), and suggest that broad patterns of assortative mating may be due in part to variation in the availability of homo- vs. heterotypic individuals as mates.

#### Reproductive isolation index

Applying an index of premating reproductive isolation (RI) allowed us to control for variation in available mates at a fine geographic scale (Figure S3). In the *rustica-tytleri* transect in Russia, both parentals and hybrids co-occurred in several populations (Figure S3A). Assortative mating by genotype was comparatively weak in these populations (Figure S3A). However, in all populations where both parental forms coexisted, there was evidence for assortative mating by migratory phenotype (average RI= 0.28). Isolation was strongest among *rustica* individuals (RI= 0.52); that is, individuals assigned *rustica* migratory phenotypes were >50% more likely to pair with each other than with a *tytleri* migratory phenotype. Assortative mating by color was less consistent among populations (Figure S3A). This result suggests a central role for divergent migratory behavior in mediating premating reproductive isolation between *rustica* and *tytleri*.

There was some assortative mating by genotype in the *rustica-gutturalis* transect in China (average RI= 0.14, Figure S3B). However, this was due to the absence of parentals from the hybrid zone center and consequent high pairing frequency among hybrids (“conspecific” matings); indeed, there was no population in which parental *rustica* and *gutturalis* co-occurred (Figure S3B). There was some weak assortative mating by migratory phenotype in the hybrid zone center (Figure S3B), but mating was otherwise random based on phenotype.

In contrast to the two migratory divides, we did not detect assortative mating across the *tytleri-gutturalis* transect in China. Genotype frequencies were fairly homogeneously admixed across the transect, and both migratory phenotype and color varied little, making the question of premating isolation less relevant (Figure S3C). Taken together, our measurement of premating barriers suggests stronger assortative mating by migratory phenotype than color in both migratory divides. However, the distributions of parental vs. hybrid genotypes, and hence potential mates, varied substantially. The mechanism by which migratory divides contribute to reproductive barriers may therefore differ between subspecies pairs (Figure S3).

## Discussion

We tested the hypothesis that migratory divides are broadly important to the maintenance of reproductive barriers between barn swallow subspecies by sampling comprehensively across multiple contact zones. Our analyses collectively suggest that 1) there was less hybridization across transects with migratory divides than across transects without migratory divides; 2) divergent migratory behavior explained large proportions of genetic variance relative to other traits within migratory divides; and 3) divergent migratory behavior *per se* contributed to premating reproductive barriers. Further, geographic coincidence between migratory divides and narrow hybrid zones supports a longstanding hypothesis (Irwin & Irwin 2005) that divergent migratory routes around the Tibetan Plateau maintain range boundaries in Siberian and central Asian avifauna.

Many birds that breed in Asia circumnavigate the inhospitable Tibetan Plateau to the east or west en route to wintering grounds in south Asia or Africa (Irwin & Irwin 2005). By sampling most of the Asian range of the barn swallow, we found multiple migratory divides centered at the same longitude (∼100 degrees) but at different latitudes and between different subspecies pairs. These narrow hybrid zones occurred across regions with no obvious ecological gradients or barriers to dispersal, suggesting isolation is not due to divergent ecological selection during the breeding season. Instead, the striking coincidence in width and geographic locations of the hybrid zones, and the similar proportions of backcrosses in each zone (Figure S2), suggest that hybrid zones have independently settled in regions where selection against hybrids is symmetrical (Price 2008) or costs of long-distance migration are minimized (Toews 2017). Such observations implicate a major barrier that drives both the location and extent of hybridization across a broad geographic region. Limited hybridization in these areas is the pattern we would predict if the Tibetan Plateau shapes differences in migratory behavior and contributes to the maintenance of species boundaries.

Social pairing data further suggest that assortative mating by migratory phenotype may be an important premating barrier to hybridization between *rustica* and *tytleri.* However, although migratory phenotype explained large proportions of genetic variance, premating isolation was weaker between *rustica* and *gutturalis* in China, likely due to the absence of parental individuals in the center of the hybrid zone. In birds, it has been proposed that premating barriers often arise early in divergence, with postmating barriers and reinforcement appearing later via selection against unfit hybrids (Price 2008). Different isolating mechanisms operating within the two migratory divides may reflect different lengths of time since secondary contact, as well as contributions of other variables, such as competitive exclusion or unmeasured ecological factors, to isolation. Intrinsic postmating barriers are unlikely given shallow divergence (Zink *et al*. 2006; Smith *et al*. 2018), presence of backcrosses in all hybrid zones, and the absence of fixed differences between any subspecies pair. It remains possible that as-yet-undetected loci are associated with divergent migratory behaviors and cause intrinsic genetic incompatibilities in hybrids. However, many other migratory divides lack evidence for hybrid unfitness or genetic differentiation associated with migratory phenotypes (Davis *et al*. 2006; Liedvogel *et al*. 2014; Ramos *et al*. 2017; Toews *et al*. 2017). It is therefore more likely that assortative mating and extrinsic selection against hybridization maintain narrow hybrid zones at migratory divides, although we cannot assess the relative importance of pre- vs. postmating barriers with our current data.

Here we present evidence for a central role of divergent migratory behavior in the maintenance of reproductive boundaries across replicated hybrid zones, supporting a longstanding but rarely evaluated hypothesis that migratory behavior can be an important engine of speciation. Future work studying hybrid fitness will further clarify the mechanisms by which reproductive isolation is maintained within migratory divides.

## Acknowledgements

We are grateful to the State Darwin Museum in Moscow, Hainan Normal University, National University of Mongolia, the Mongolian Ornithological Society, and the Japan Bird Research Association for sponsoring our research. Work was conducted in accordance with University of Colorado IACUC protocol #1303. Many people assisted with fieldwork, including Caroline Glidden, Rachel Lock, Yulia Sheina, Nikolai Markov, Gennady Bachurin, Olga Zayatseva, Elena Shnayder, Unurjargal Enkhbat, Bayanmunkh Dashnyam, Davaadorj Enkhbayar, Wataru Kitamura, Takashi Tanioka, and Yuta Inaguma. Brittany Jenkins prepared the genomic libraries. Dai Shizuka advised on the phenotype networks. We thank Amanda Hund, Bruce Lyon, Trevor Price, Scott Taylor, and the Safran and Taylor labs at the University of Colorado for feedback on the manuscript. This project was funded by NSF-CAREER grant DEB-1149942 to RJS and the National Geographic Society Committee on Research and Exploration grant to ESCS.

