## Supplemental figures and methods for "Migratory divides coincide with species barriers across replicated avian hybrid zones above the Tibetan Plateau"

**Figure S1:** Phenotypic variation in each parental subspecies and in the three hybrid zones. Colors correspond to geographic sampling regions, as in the map in Figure 2. A) there are significant differences in carbon isotope ratio between *rustica –tytleri* (red-gold) and *rustica- gutturalis* (red-blue), but not between *tytleri* and *gutturalis* (gold-blue). Wing length (B) and breast chroma (C) differ significantly among the parental subspecies (all $p < 0.001$). Throat chroma (D) is significantly darker and tail length (E) significantly shorter in *gutturalis,* but neither trait differs between *rustica* and *tytleri. Rustica* has significantly more pointed wings (F) than *tytleri* and *gutturalis*. Genomic data show that swallows in eastern Russia and northeastern China, currently taxonomically classified as "*gutturalis*", consistently had 20-25% *tytleri* ancestry, as did nearly all individuals captured in Japan. These birds were frequently phenotypically intermediate between *tytleri* and *gutturalis*, as indicated by the on-average darker and more *tytleri*-like color of TG hybrids (panel C). Most birds captured in eastern Mongolia had between 40 and 90% *tytleri* ancestry, with a few parental *tytleri* occurring in central Mongolia.

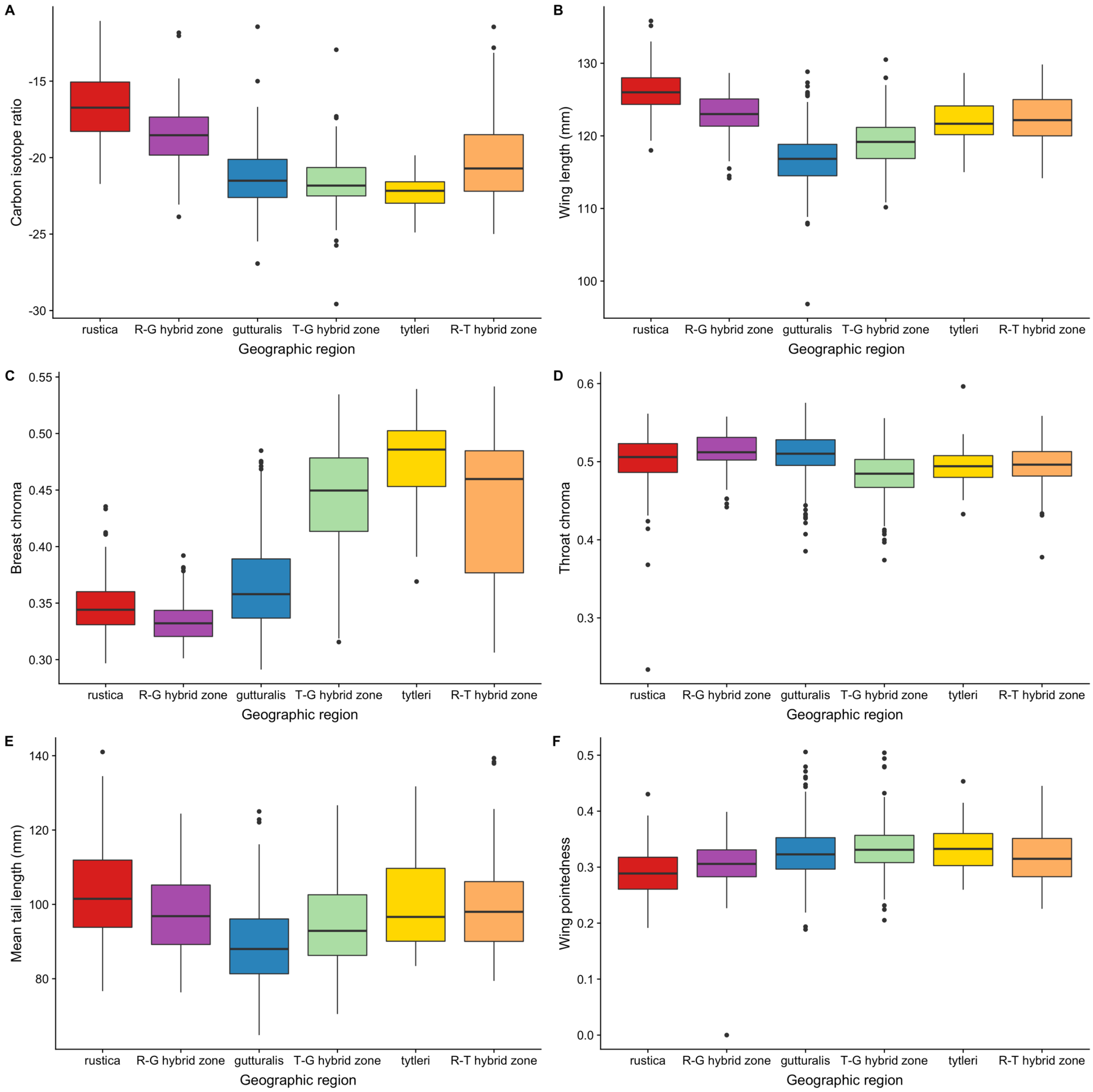

**Figure S2:** Hybrid class assignment for each pair of subspecies. F1 and early generation hybrids have intermediate hybrid indices and high average heterozygosity, while backcrosses have hybrid indices closer to 0 or 1 and lower average heterozygosity. Parental individuals have a hybrid index of 0 or 1 and low heterozygosity. Note that there are comparatively few early generation hybrids in the *rustica-gutturalis* (left panel) and *rustica-tytleri* (center panel) hybrid zones, indicating fairly strong reproductive isolation, whereas there are many F1 and later generation hybrids between *tytleri* and *gutturalis* (right panel).

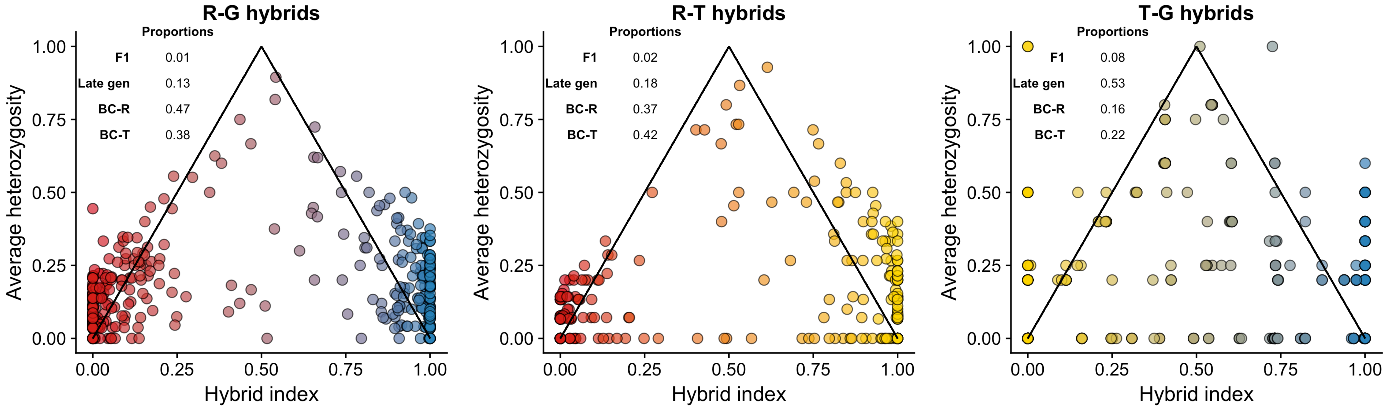

**Figure S3**: Strength of reproductive isolation (RI) based on ancestry, migratory phenotype and ventral color in A) the *rustica-tytleri* transect in Siberia, B) the *rustica-gutturalis* transect in China, and C) the *tytleri-gutturalis* transect in China. The top row of each plot shows the strength of RI in each population along the transect, with each point corresponding to the RI calculated for a particular pair type (e.g. *rustica-rustica* pairs, *rustica-tytleri* hybrid pairs, etc). Pair type colors are shown in side legend. The bottom panel shows the corresponding fastSTRUCTURE plots of the distribution of genotypes in each population. The RI index ranges from -1 to 1, with an RI index of 0 (dashed line) indicating random mating with respect to a particular trait. An RI index > 0 (above the dashed line) indicates assortative mating, and an RI index below the dashed line indicates disassortative mating. Points are jittered slightly to enhance plot readability. Points at the very bottom of the plot (below the dotted line) indicate pairings that randomization tests predicted to occur in the population based on the distribution of genotypes and phenotypes from all individuals, but were not observed in the subset of individuals for which we had pairing data. Note that in all three transects there are “unexpected” hybrid genotypes despite our cutoff that required hybrids have >20% assignment probability to each of two clusters. There is a single *tytleri-gutturalis* individual in the “*rustica-tytleri*” transect in Russia (panel A, population 3). There are *tytleri-gutturalis* hybrids at the eastern edge of the “*rustica-gutturalis*” transect in China (panel B, populations 6-8) and a single *rustica-tytleri* hybrid in the western part of that transect (population 4). Likewise, there are *rustica-gutturalis* hybrids across much of the “*tytleri-gutturalis*” transect in China (panel C). A) There was weak assortative mating by genotype in the *rustica-tytleri* transect, but stronger assortative mating by migratory phenotype and, to a lesser extent, ventral coloration. B) There was assortative mating by ancestry in the *rustica-gutturalis* transect due to the predominance of hybrids in the center of the hybrid zone, but weak assortative mating by phenotype. C) Fairly homogeneous admixture within populations across the *tytleri-gutturalis* transect resulted in little to no premating isolation in this transect.

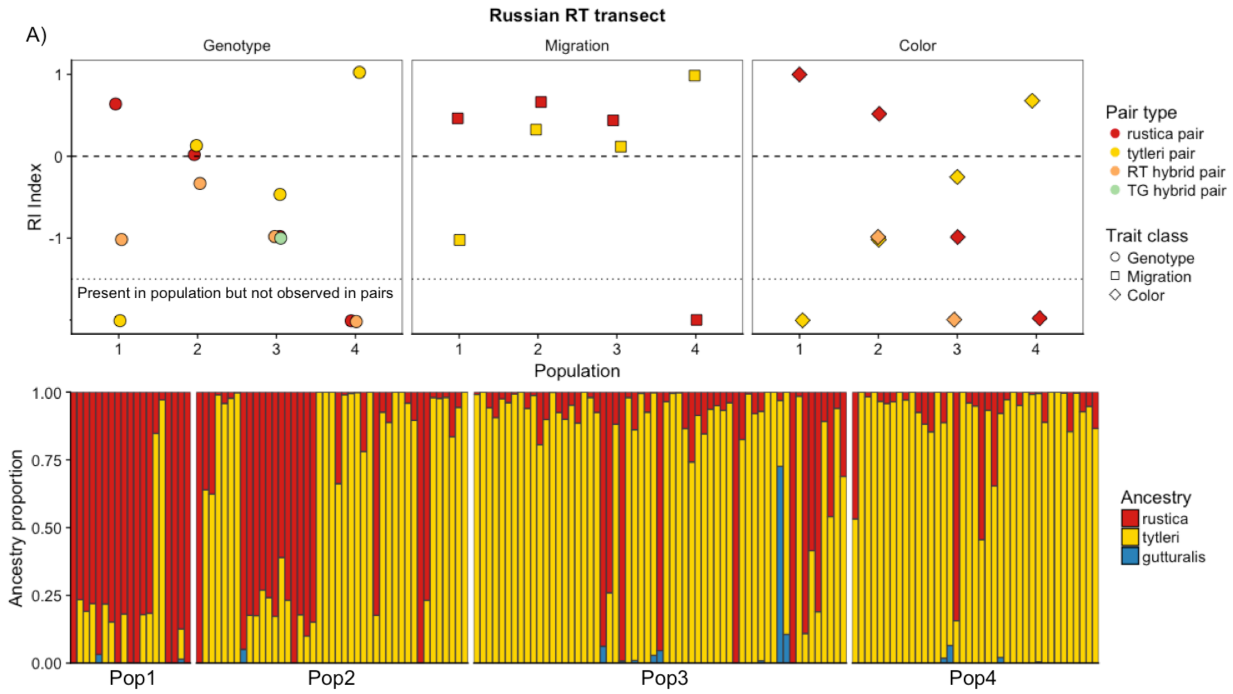

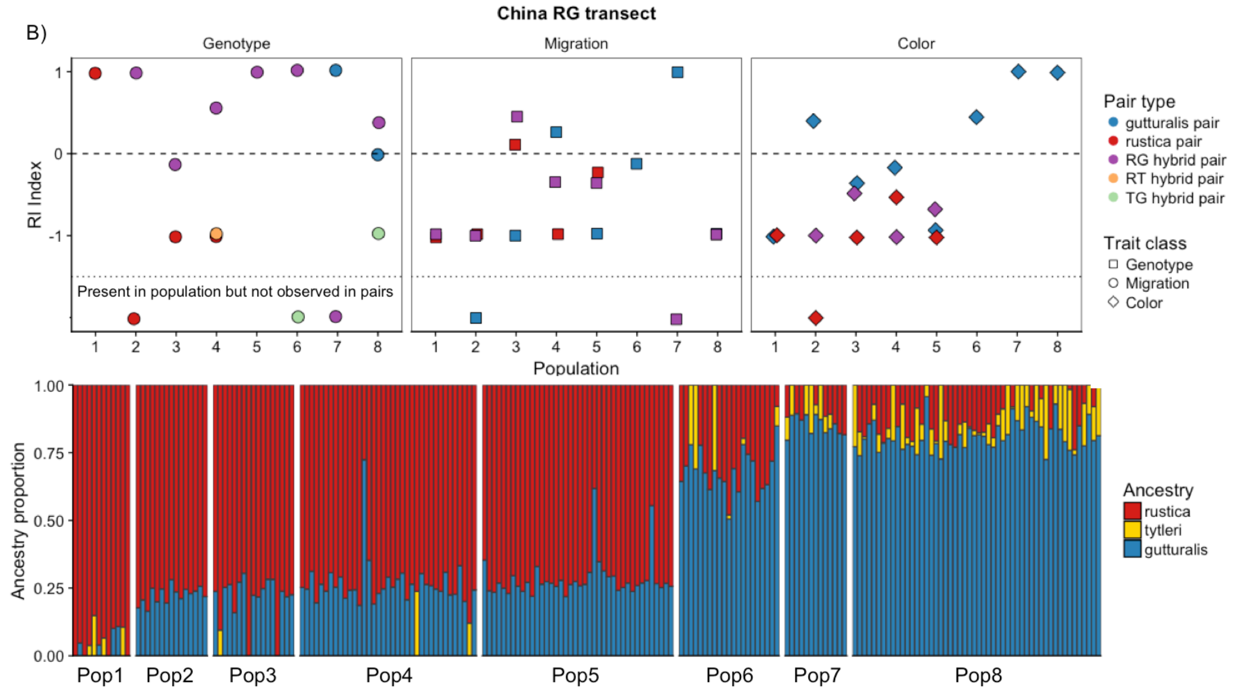
China RG transect
B)
Genotype
Migration
Color
RI Index
Present in population but not observed in pairs
Population
Pair type
gutturalis pair
rustica pair
RG hybrid pair
RT hybrid pair
TG hybrid pair
Trait class
Genotype
Migration
Color
Ancestry proportion
Ancestry
rustica
tytleri
gutturalis
Pop1
Pop2
Pop3
Pop4
Pop5
Pop6
Pop7
Pop8

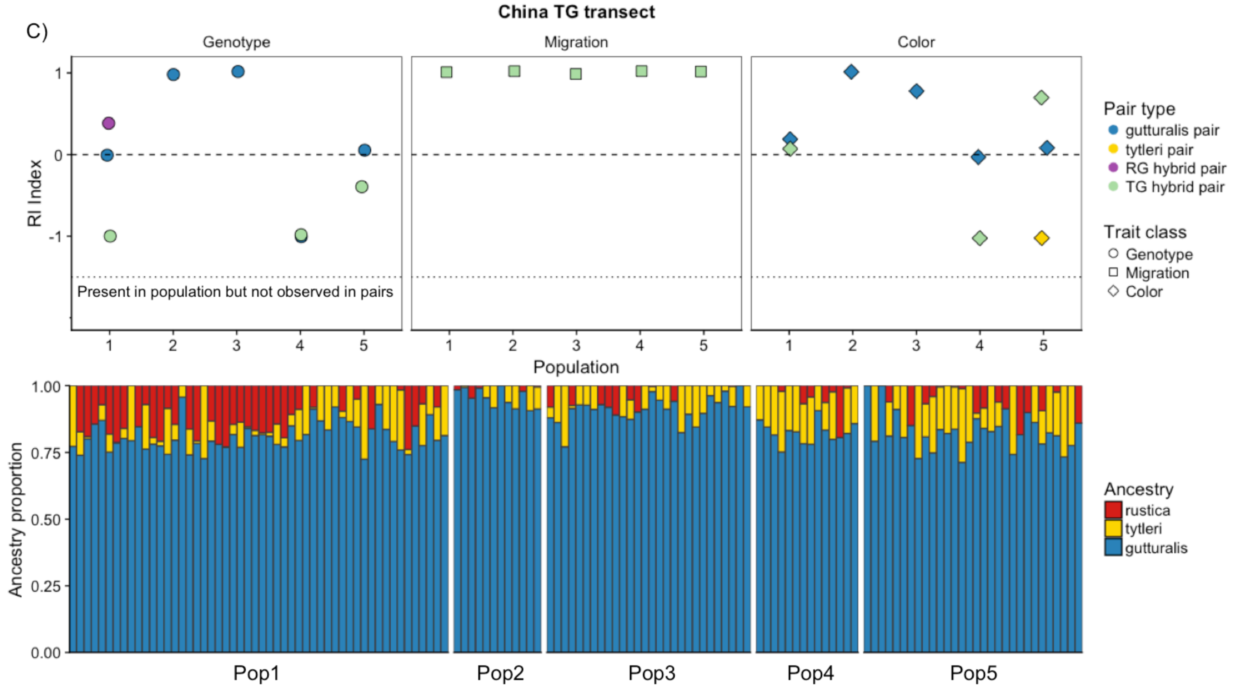
China TG transect
C)
Genotype
Migration
Color
RI Index
Present in population but not observed in pairs
Population
Pair type
gutturalis pair
tytleri pair
RG hybrid pair
TG hybrid pair
Trait class
Genotype
Migration
Color
Ancestry proportion
Ancestry
rustica
tytleri
gutturalis
Pop1
Pop2
Pop3
Pop4
Pop5

**Table S1:** Geographic clines for non-focal morphometric traits. Boldfaced clines are those that have centers coincident with the ancestry cline (Table 1 in main text). Starred widths are narrower than expected under a neutral diffusion model assuming a dispersal distance of 42 km and a hybrid zone age older than 20 years. Italicized clines show no statistically significant variation in trait values across the transect (ANOVAs followed by Benjamini-Hochberg correction for multiple testing) and are consequently poorly described by cline models. The § symbol shows clines that are wider than expected with a dispersal distance of 42km, but narrower than expected if dispersal is 100km and clines are older than 20 years. Clines centers are in kilometers from the westernmost transect point.

| Transect | Wing length center | Wing length width (km) | Wing pointedness center | Wing pointedness width (km) | Wing convexity center | Wing convexity width (km) | Tail length center | Tail length width (km) | Throat chroma center | Throat chroma width (km) |
|---|---|---|---|---|---|---|---|---|---|---|
| RG-China | **1183.33 (945.67-1994.54)** | 1173.12 (585.47-2029.99) | *1445.18 (1323.88-1512.46)* | *7.64 (0.54-168.05)* | *981 (600.35-1671.81)* | *120.48 (3.99-2139.56)* | *823.47 (347.12-868.48)* | *6.39 (0-171.1)* | 297.76 (77.81-3.12) | 712.59 (0-244.61) |
| RT-Mongolia | **395.14 (-28.76-1367.38)** | 730.68 (0.08-2029.97) | **592.58 (99.82-823.79)** | 91.56* (0.02-2142.06) | *563.64 (343.35-2199.98)* | *79.08 (0.01-242.57)* | *1775.66 (1145.54-2200)* | *88.29 (0-1009.34)* | *1323.78 (69.21-1147.74)* | *2199.99 (1.25-549.45)* |
| RT-Russia | **787.69 (776.3-797.61)** | 2.84 * (0-82.41) | **766.77.82 (745.92-787.93)** | 2.82* (0-106.01) | *557.88 (79.93-957.89)* | *46.24 (0.49-2158.11)* | *1967.05 (1401.77-2200)* | *67.07 (0-606.43)* | *2001.57 (237.81-13.35)* | *2200 (0-2229.42)* |
| TG-China | **565.77 (2.09-891.8)** | 861.66 (61.46-2029.97) | *1866.13 (975.67-2200)* | *140.82 (0-1069.19)* | *772.47 (689.31-976.48)* | *15.55 (0.58-455.17)* | **630.46 (401.52-881.53)** | 107.41* (0.01-788.69) | *797.85 (38.01-494.88)* | *797.85 (38.01-203.97)* |
| TG-Mongolia | *1018.8 (636.94-1273.81)* | *1088.94 (3.61-2028.58)* | *889.77 (594.88-2200)* | *56.02 (0.01-2108.98)* | *2156.15 (594.75-2199.99)* | *965.14 (0-1426.69)* | *22.62 (0-1207.31)* | *684.16 (595.22-2199.97)* | *-27.76 (211.15-30)* | *210.58 (0.05-1447.12)* |
| TG-Russia | **621.26 (188.05-712.33)** | 597.37 (284.12-2029.93) | *678.16 (602.94-903.60)* | *17.38 (0.02-71.35)* | *244.05 (4.66-405.88)* | *964.02 (1.19-2167.75)* | *307.12 (-29.98-1334.23)* | *1670.83 (5.18-2229.98)* | *254.07 (666.32-28.26)* | *526.47 (489.17-2228.01)* |

**Table S2:** Results of redundancy analysis for each transect. The first model for each transect ("all traits") shows genetic variance partitioned among all sets of traits (color, geography, and migratory phenotype). The subsequent models show genetic variance partitioned among each set of traits individually, in a model that is conditioned on the other traits.

| Transect | Model | F | P Value |
|---|---|---|---|
| RG- China | All traits$_{4,146}$ | 111 | **0.001**** |
| | Color$_{1,146}$ | 2.67 | 0.106 |
| | Geography$_{2,146}$ | 120.24 | **0.001**** |
| | Migration$_{1,146}$ | 13.54 | **0.001**** |
| RT- Russia | All traits$_{4,109}$ | 45.22 | **0.001**** |
| | Color$_{1,109}$ | 51.76 | **0.001**** |
| | Geography$_{2,109}$ | 3.24 | 0.058 |
| | Migration$_{1,109}$ | 7.39 | **0.008**** |
| RT-Mongolia | All traits$_{4,34}$ | 16.33 | **0.001**** |
| | Color$_{1,34}$ | 6.5 | **0.014*** |
| | Geography$_{2,34}$ | 9.97 | **0.001**** |
| | Migration$_{1,34}$ | 0 | 0.948 |
| TG- China | All traits$_{4,46}$ | 13 | **0.001**** |
| | Color$_{1,46}$ | 0.19 | 0.63 |
| | Geography$_{2,46}$ | 20.68 | **0.001**** |
| | Migration$_{1,46}$ | 1.3 | 0.273 |
| TG- Mongolia | All traits$_{4,62}$ | 7.73 | **0.001**** |
| | Color$_{1,62}$ | 26.1 | **0.001**** |
| | Geography$_{2,62}$ | 1.04 | 0.346 |
| | Migration$_{1,62}$ | 0.04 | 0.827 |
| TG- Russia | All traits$_{4,69}$ | 17.43 | **0.001**** |
| | Color$_{1,69}$ | 7.63 | **0.005**** |
| | Geography$_{2,69}$ | 21.18 | **0.001**** |
| | Migration$_{1,69}$ | 0.01 | 0.898 |

### Table S3: Sampling locations and sample sizes

| Country | Location | Subspecies | Latitude | Longitude | N |
|---|---|---|---|---|---|
| China | Urumqi | *rustica* | 43.817486 | 87.627925 | 12 |
| China | Dunhuang | *rustica* | 40.093024 | 94.670341 | 15 |
| China | Golmud | *rustica* | 36.43766325 | 94.76997025 | 4 |
| China | Yumen | RG hybrids | 40.28389789 | 97.03055778 | 18 |
| China | Jiuquan | RG hybrids | 39.7414906 | 98.51988518 | 40 |
| China | Gaotai | RG hybrids | 39.361411 | 99.813419 | 4 |
| China | Zhangye | RG hybrids | 38.93521733 | 100.4403971 | 36 |
| China | Wuwei | RG hybrids | 37.888956 | 102.626964 | 21 |
| China | Lanzhou | *gutturalis* | 36.08753921 | 103.7644974 | 14 |
| China | Yinchaun | *gutturalis* | 38.52959363 | 106.1747908 | 30 |
| China | Nanning | *gutturalis* | 22.79191794 | 108.3262965 | 17 |
| China | Xian | *gutturalis* | 34.34565427 | 108.8044669 | 22 |
| China | Hainan | *gutturalis* | 19.2264018 | 109.146506 | 20 |
| China | Baotu | *gutturalis* | 40.55582573 | 110.003829 | 26 |
| China | Changsha | *gutturalis* | 28.40071539 | 112.8074179 | 18 |
| China | Zhengzhou | *gutturalis* | 34.82433361 | 113.6707127 | 28 |
| China | Beijing | *gutturalis* | 39.81955823 | 116.3329121 | 13 |
| China | Qinhuangdao | *gutturalis* | 39.92558 | 119.594752 | 28 |
| China | Shenyang | *gutturalis* | 41.80877479 | 123.5269676 | 14 |
| China | Qiqihar | *gutturalis* | 47.34003877 | 123.9758888 | 26 |
| China | Changchun | *gutturalis* | 43.89488661 | 125.320679 | 31 |
| China | Harbin | *gutturalis* | 45.761237 | 126.609063 | 43 |
| China | Shuangyashan | *gutturalis* | 46.65299473 | 131.1480528 | 33 |
| Japan | Tokyo | *gutturalis* | 35.95222638 | 139.1258438 | 29 |
| Japan | Hokkaido | *gutturalis* | 42.309632 | 142.525039 | 15 |
| Mongolia | Khovd | *rustica* | 48.009753 | 91.661555 | 8 |
| Mongolia | Durgun | *rustica* | 48.33141673 | 92.63146527 | 11 |
| Mongolia | Urgamal | *rustica* | 48.515155 | 94.2987396 | 5 |
| Mongolia | Santmorgaz | *rustica* | 48.584554 | 95.433157 | 7 |
| Mongolia | Tsetserleg | RT hybrids | 47.460351 | 101.474926 | 1 |
| Mongolia | Ugii Lake | TG hybrids | 47.7509 | 102.772322 | 6 |
| Mongolia | Kharkhonh | TG hybrids | 47.191711 | 102.856199 | 16 |
| Mongolia | Dashinchilin | TG hybrids | 47.845526 | 104.0568 | 12 |
| Mongolia | Lun | TG hybrids | 47.866496 | 105.211692 | 14 |
| Mongolia | Zuunmod | TG hybrids | 47.698606 | 106.95476 | 10 |
| Mongolia | Cincer Erdene/ Tsenhermandel | TG hybrids | 47.71765055 | 108.8470845 | 11 |
| Mongolia | Chingis Khan | TG hybrids | 47.331896 | 110.686229 | 20 |
| Mongolia | Berh | TG hybrids | 47.783939 | 111.165747 | 6 |
| Mongolia | Batnorov/Norovlin | TG hybrids | 48.44214767 | 111.8312897 | 6 |
| Mongolia | Bayan Ovoo Bridge | TG hybrids | 47.763829 | 112.164689 | 7 |
| Mongolia | Bayan.Uul | TG hybrids | 49.122033 | 112.677635 | 15 |
| Mongolia | Holonbur/Norovlin | TG hybrids | 48.469793 | 113.1130633 | 12 |
| Mongolia | Bulgan.Soum.Dornod | TG hybrids | 48.000609 | 113.936327 | 14 |

| | | | | | |
|---|---|---|---|---|---|
| Mongolia | Dashbalbar | TG hybrids | 49.5092496 | 114.4579721 | 20 |
| Russia | Moscow | *rustica* | 56.76593274 | 37.78463094 | 31 |
| Russia | Yekaterinburg | *rustica* | 57.54072522 | 62.71656765 | 51 |
| Russia | Karasuk | *rustica* | 53.93487526 | 77.73869885 | 27 |
| Russia | Novosibirsk/ Krasny Yar | *rustica* | 55.7253224 | 86.1515802 | 10 |
| Russia | Krasnoyarsk/Kansk | *rustica* | 56.35105281 | 93.05357331 | 16 |
| Russia | Kontorskaya | *rustica* | 56.02804313 | 97.86840525 | 8 |
| Russia | Aculshet/ Berezovka/ Byronovka | RT hybrids | 55.87410464 | 98.06904573 | 11 |
| Russia | Oblepiha | RT hybrids | 55.667468 | 98.448748 | 8 |
| Russia | Alzamay | RT hybrids | 55.59150071 | 98.609405 | 14 |
| Russia | Zamzor | RT hybrids | 55.3732061 | 98.65219881 | 21 |
| Russia | Mara | RT hybrids | 55.00063873 | 98.845692 | 15 |
| Russia | Kamenka | RT hybrids | 55.00048 | 98.848568 | 20 |
| Russia | Uk | RT hybrids | 55.07751313 | 98.87160958 | 24 |
| Russia | Hingui Station | RT hybrids | 54.7937232 | 99.3445835 | 10 |
| Russia | Hingui Village | RT hybrids | 54.798431 | 99.440598 | 20 |
| Russia | Umigan | *tytleri* | 54.66629833 | 100.0836122 | 9 |
| Russia | Male-Kutulyk | *tytleri* | 53.40102078 | 102.7856412 | 9 |
| Russia | Malamolevo | *tytleri* | 53.39322367 | 102.862147 | 6 |
| Russia | Zakaltoose | *tytleri* | 52.021259 | 106.590942 | 30 |
| Russia | Nikolaevska | TG hybrids | 51.06395 | 111.791206 | 10 |
| Russia | Tataurova | TG hybrids | 51.607376 | 112.939376 | 7 |
| Russia | Narasun | TG hybrids | 50.08136 | 112.970703 | 14 |
| Russia | Mixed Barns | TG hybrids | 50.46408345 | 113.4537446 | 29 |
| Russia | Narin-Talacha | TG hybrids | 51.93720487 | 114.9683183 | 45 |
| Russia | Mogocha | TG hybrids | 53.720489 | 119.781143 | 8 |
| Russia | Daktuy/ Civaky | *gutturalis* | 52.83969914 | 126.5893513 | 7 |
| Russia | Vozhaevka | *gutturalis* | 50.741612 | 128.72934 | 15 |
| Russia | Kundor | *gutturalis* | 49.103321 | 130.758652 | 9 |
| Russia | Yadrina | *gutturalis* | 48.969479 | 131.018448 | 11 |
| Russia | Chernaevka | *gutturalis* | 44.332958 | 132.517776 | 17 |
| Russia | Richnoy | *gutturalis* | 45.94273 | 133.882263 | 10 |
| Russia | Magelevka | *gutturalis* | 47.968128 | 134.912735 | 16 |

**Supplemental methods**

*Quantification of plumage color*

Body feathers collected were mounted on white index cards (Safran & McGraw 2004) and color was quantified using a fiber optic cable attached to an Ocean Optics USB-4000 spectrophotometer and an Ocean Optics PX-2 pulsed xenon light source. Each patch (throat, breast, belly, and vent) was measured 3 times, with 20 scans averaged per measurement. The three measurements per patch were then averaged together, and we calculated the hue, chroma, and brightness for each color patch for each individual bird (Safran *et al.* 2010). We conducted a principal components analysis (PCA) on the correlation matrix of each of the three color measures (hue, saturation, brightness) for each of the four patches (throat, breast, belly, and vent). Breast chroma loaded strongly on PC1 and throat chroma loaded strongly on PC2. We consequently used breast chroma as the measure of ventral color in our analyses and throat chroma as our measure of throat color.

*Stable Isotope Analysis*

We determined variation in wintering grounds by analyzing stable isotope ratios in the inner-most tail rectrices. We focused on carbon isotopes, as variation in this element corresponds to differences in the distribution of C3 and C4 plants (Chamberlain *et al.* 2000). Previous records of barn swallow migratory routes suggest that *rustica* overwinters in drier regions of Africa and *gutturalis* and *tytleri* overwinter in the wetter regions of south and southeast Asia. We cleaned tail feathers in a 2:1 mix of chloroform and methanol to remove dirt and oils. We then cut 1mg of feather vanes from the center of the feather, avoiding the feather tip and base. Feather tissue was rolled into 4 x 6 mm tin capsules (Costech Analytical Technologies, Valencia, CA) and analyzed at the U.S. Geological Survey Stable Isotope Laboratory (Denver, CO). Samples were combusted in an elemental analyzer (Carlo Erba NC2500, Milan, Italy) interfaced to a Micromass Optima mass spectrometer (Fry *et al.* 1992). Data are reported in standard delta notation with respect to internationally accepted scales (Air and V-PDB) following normalization with USGS 40 ($\delta^{15}N$ = -4.52 ‰, $\delta^{13}C$ = -26.24 ‰) and 41 ($\delta^{15}N$ = 47.57 ‰, $\delta^{13}C$ = 37.76 ‰). Analytical precision of replicate standards, including two additional secondary standards was better than +/- 0.2 ‰ for both isotopes.

*Genotyping- by- sequencing and variant calling*

We extracted genomic DNA from blood samples using DNEasy blood and tissue kits (Qiagen) following a version of the standard protocol modified to include an overnight digestion. Libraries for RAD-sequencing were prepared following Parchman *et al.* (2012). Briefly, we digested genomic DNA with the restriction enzymes MseI and EcoRI. A unique 8, 9, or 10-base barcode was ligated to the fragment libraries for each individual. We then pooled barcoded samples and amplified fragments using standard Illumina primers. Libraries were size-selected (350-400 base pairs) using a PippinPrep quantitative electrophoresis unit (Sage Science). We sequenced 100bp single-end reads on an Illumina HiSeq 2500 and HiSeq4000 at the University of Texas, Austin genomic sequencing and analysis facility. We ran three different libraries on four replicate lanes (12 lanes total). Samples from Russia were run in 2013 with 538 samples multiplexed per lane; samples from China, Mongolia, and Japan were run with 536 multiplexed samples in 2015, and additional samples from China were run with 546 multiplexed samples in 2016. The 2016 library included 47 samples from the two previous libraries to control for lane and year effects. We ran roughly the same number of samples per lane for each library to ensure consistency in read depth per individual. We found no lane effects in samples run in multiple years.

After sequencing, reads were filtered for contaminant sequences (*E. coli*, *Phi X* and remnant Illumina oligos) and bases associated with the restriction enzyme cut sites were removed. The base following the restriction cut site was lower quality on average and was also removed. Trimmomatic (Bolger *et al.* 2014) was used to remove bases with quality below 30 from the ends of each read, and reads with average quality below 30 or length below 50bp were removed. Individuals with < 100,000 reads (n=6) and were discarded from further analysis. This left 1288 birds in the final dataset. Reads were aligned to the barn swallow genome using bwa mem, and SNPs were identified using *bcftools* and *samtools* (Li & Durbin 2009; Li *et al.* 2009). To ensure only high-quality SNPs were retained for analysis, we required a median read depth of 7 reads per locus and 80% of individuals to have data at that locus. We used a minor allele frequency cutoff of 5% to filter out rare variants and included only one SNP per 100bp to remove tightly linked loci and minimize inclusion of sequencing errors. This resulted in 12,383 SNPs. To improve precision in subsequent analyses, we incorporated genotype uncertainty (from PL fields of the *samtools* output) into our final genotype scores, resulting in continuous genotype probabilities for each individual at each locus ranging from 0 to 2, with 0 and 2 as the homozygotes and 1 as the heterozygote. For analyses that required fixed genotypes, we rounded genotype probabilities as follows: 0 = 0-0.1; 0.95-1.05 = 1; 1.9-2.0= 2. All other genotypes were scored as NA. This rounding is conservative, and strong agreement between analyses of population structure that employ continuous genotypes and those that use rounded genotypes suggest the rounding is appropriate.

*Assignment to hybrid classes*

To determine whether gene flow is ongoing (presence of F1 and backcrossed individuals) between the different subspecies, we assigned individuals to hybrid classes. All methods of hybrid class assignment assume hybridization between only two differentiated populations. We therefore replicated this analysis for each of the three subspecies pairs. Because fastSTRUCTURE returned higher assignment probabilities to individual clusters than TESS, we used fastSTRUCTURE assignments in the following analyses. We defined birds assigned with >90% probability to a single cluster as “parentals.” Birds with >10% of their genome assigned to two clusters were designated as two-way hybrids. Individuals with >10% assignment probability to a third cluster (n=25) were dropped from these analyses. We also dropped birds with <90% assignment to a parental cluster and <10% assignment to additional clusters (e.g. individuals with 84% to one cluster and 8% to two additional clusters, n=19) as it was not possible to determine the direction of hybridization. Parental assignments in fastSTRUCTURE agreed with our classifications based on phenotypes in the field, with the majority of *rustica* parentals occurring in the western regions of Russia, China, and Mongolia, parental *tytleri* occurring around Lake Baikal and in central Mongolia, and parental *gutturalis* occurring in southern China (Figure 1, 2).

We calculated $F_{ST}$ between parental pairs (*rustica-tytleri*, *tytleri-gutturalis, rustica-gutturalis*) in the R package outflank (Lotterhos & Whitlock 2015). Average pairwise $F_{ST}$ was 0.028 between *rustica* and *gutturalis*, 0.019 between *tytleri* and *gutturalis,* and 0.016 between *rustica* and *tytleri*. We used the outFLANK results to identify differentiated markers informative for assigning non-parental individuals to hybrid classes. There were no diagnostic loci ($F_{ST}$ =1) segregating between any pair of subspecies, so we defined ancestry- informative markers as loci with $F_{ST}$ > 0.41 between a pair. This cutoff corresponded to the top 0.5% of loci from the shallowest comparison (*tytleri* - *gutturalis*), and produced the minimum number of loci (n=5) needed for hybrid class assignment (Gompert & Buerkle 2009). Our $F_{ST}$ cutoff yielded 29 differentiated loci between *rustica-gutturalis*, 15 loci between *rustica-tytleri*, and 5 loci between *tytleri* and *gutturalis.* The small number of differentiated loci between *tytleri* and *gutturalis* means that specific hybrid class assignments should be interpreted with caution in this pair.

We used the R package *introgress* (Gompert & Buerkle 2009) to calculate maximum likelihood estimates of hybrid index (i.e. the proportion of alleles inherited from each parent) and average heterozygosity for individuals identified as two-way hybrids. This method makes few assumptions about selection and linkage, and is therefore appropriate for the barn swallow system, where divergence is shallow and differentiated loci may be under selection (Scordato *et* *al.* 2017). If there are fixed differences between parentals, F1s are expected to have a hybrid index = 0.5 and average heterozygosity = 1, whereas later generation hybrids and backcrosses will have lower average heterozygosity and intermediate hybrid indices. However, assignment to hybrid classes in comparisons without diagnostic loci is inexact (Buerkle 2005), particularly when there are few differentiated loci with missing data. We consequently followed recommendations of recent studies (Milne & Abbott 2008; Bouchemousse *et al.* 2016; Walsh *et al.* 2016) and broadly grouped individuals into four different classes. Individuals with hybrid indices ranging from 0.02-0.25 or 0.75-0.98 were considered backcrosses to one or the other parental type. Individuals with intermediate hybrid indices (>0.25 and <0.75) and heterozygosity ≥0.80 were classified as F1 hybrids. Individuals with intermediate hybrid indices but lower average heterozygosity (<0.80) were classified as later generation hybrids. Although these categories are broad, they are appropriate for determining whether there is ongoing gene flow across hybrid zones. Differentiating F1 from later generation hybrids was particularly challenging for the *tytleri-gutturalis* comparisons given the very small number of differentiated loci, and we therefore interpret these assignments with caution.

*Geographic cline analysis*

We used geographic clines to quantify changes in the frequencies of genotypes and phenotypes across transects through three narrow hybrid zones (*rustica-tytleri*, *rustica-gutturalis*) and three areas of extensive admixture (*tytleri-gutturalis*). We used PC1 as a measure of ancestry rather than cluster assignment probabilities from fastSTRUCTURE or TESS because it is a single measure that captures the continuous gradient of variation we observed in ancestry across the three parental groups. We could then fit clines to the same, single measure of ancestry across each contact zone. We also fit clines to breast chroma, throat chroma, tail streamer length, carbon isotope value, wing convexity, wing pointedness, and wing length. We normalized all traits and the genetic PC scores to values between 0 and 1 using the formula

$$z_i = \frac{x_i - \min(x)}{\max(x) - \min(x)}$$

Where $x$ is the vector of trait values and $z_i$ is the normalized trait value for individual $x_i$.

Because the scale of sampling and sample sizes were variable among transects, we defined populations along 20km intervals; that is, birds sampled within 20km were grouped into a single “population.” This distance ensured that there were at least 8 birds per “population”, and that birds within the same “population” could reasonably interact with each other (Turner 2010). We calculated mean and variance for each trait in each population, and calculated distances (km) between populations using the Haversine great circle distance. We then fit clines along each sampling transect using the Metropolis-Hastings Markov chain Monte Carlo algorithm implemented in the R package HZAR (Derryberry *et al.* 2014). We fit 5 cline models with different exponential tails (none, both tails, right, left, and mirrored) to each trait and ancestry (genomic PC1) across each transect. Each cline model estimated the center (*c,* km from the westernmost transect point) and width (*w*, 1/maximum slope) of the cline, and estimated the mean and variance as free parameters. Models were initialized with $10^5$ generations of burn-in and $10^6$ chains and run with three independent MCMC chains (chain length = $10^6$). Each chain was checked for convergence and chains for each model were concatenated for model selection. We compared the five models for each trait using AICc (Cavanaugh 1997), and determined the maximum likelihood parameters for the best-fitting model. We assessed concordance between clines based on the +/- 2-log-likelihood support intervals from the best-fit model for each trait. Clines were considered concordant if the support intervals overlapped (McEntee *et al.* 2016). Cline centers with non-overlapping support intervals were interpreted to occur in different geographic locations.

We applied neutral diffusion equations to determine whether cline widths for ancestry and phenotype were narrower than expected under a scenario of no selection or reproductive isolation. Neutral variation across a contact zone is predicted to be approximately proportional to the root mean square dispersal distance per generation ($\sigma$) and the number of generations since contact (t) ($w \approx \sigma t^{0.5}$) (Barton & Gale 1993). Mean natal dispersal in British barn swallows has been estimated at 14 $\pm$ 28km (Paradis *et al.* 1998), although dispersal distances of several hundred kilometers are commonly reported (Møller 1994). Due to the uncertainty of dispersal estimates and unknown time since secondary contact, we evaluated two different dispersal distances: a conservative 42km and a less conservative 100km, assuming a one year generation time (Zink *et al.* 2006) and hybrid zone age of at least 20 years. We used these dispersal distances to calculate cline width expected under a neutral scenario for each trait along each transect. Cline widths narrower than the neutral expectation may be maintained by selection and contribute to reproductive barriers (Ruegg 2008; Brelsford & Irwin 2009).

*Assortative mating*

We used a standardized metric (Sobel and Chen 2014) to calculate the strength of premating reproductive isolation (RI) in each population along three of the transects for which we had sufficient pairing data. This RI index has primarily been used in systems where conspecifics and heterospecifics are clearly identifiable (Martin & Mendelson 2016; Lackey & Boughman 2017), and is challenging to apply to natural systems in which there is continuous variation between parental groups. We therefore defined four different genetic ancestry cutoffs for assigning individuals to “parental” or “hybrid” categories based on fastSTRUCTURE assignment probabilities. Individuals were defined as “parentals” if they had assignment probabilities to a single cluster of either 90%, 85%, 80%, or 75%. All intermediates (i.e. those with lower assignment probabilities than the cutoff) were classified as hybrids between the two parental clusters to which they had the highest assignment probabilities. We then calculated RI for each parental group in each population, as well as RI among hybrids (i.e. hybrid-hybrid pairings). We used the same population designations as in the geographic cline analyses (i.e. birds within

20km of each other were considered potential mates belonging to the same "population"). We present results from classifications based on 80% assignment probabilities; that is, each individual classified as a parental could have up to 20% ancestry from a different subspecies. Using this cutoff removed many individuals assigned as "hybrids" due to fairly small proportions of admixture, while preserving the broad patterns of hybridization observed in the geographic cline analyses.

Because mate choice is based on phenotypes rather than genotypes, we also calculated RI based on phenotype. We classified each individual to a phenotype cluster using two different sets of traits: ventral color and migratory phenotype (carbon isotope value). We assigned birds as either parentals (*rustica, tytleri,* or *gutturalis*) or hybrids (*rustica-gutturalis, rustica-tytleri, tytleri-gutturalis*) using linear discriminant analysis (LDA). For each transect, we trained a linear discriminant model on the phenotypes of 50% of the individuals in the dataset with "known" assignments to a cluster (defined based on their fastSTRUCTURE assignment probability). This training model was then used to assign the remaining 50% of individuals to a phenotype cluster. This procedure was replicated 1000 times, so that each bird had ~500 phenotypic cluster assignments. The cluster to which a bird was assigned with the highest frequency was used as its final classification. In cases where birds were assigned with approximately equal frequency to two clusters (e.g. >200 assignments as a *rustica* parent and >200 assignments as *rustica-gutturalis* hybrid), they were assigned to the cluster consistent with their genotype (n=12).

This approach resulted in each bird having three assignments as either a parental or a hybrid: an assignment based on its genotype (fastSTRUCTURE assignment probability), an assignment based on its migratory phenotype (based on ~500 replicate LDA assignments), and an assignment based on its color (based on ~500 replicate LDA assignments). We calculated the strength of isolation based on each of these three categories in each population. For example, the strength of isolation based on migratory phenotype for *rustica* in a given population is derived from the number of social pairs where both the male and female had “*rustica*” migratory phenotypes vs the number of pairs where the male and female had different migratory phenotypes. We used the same randomization method as for the genotypes to generate expected pairings based on phenotype distributions within a population, and weighted our estimates of RI by these expectations.
